# RNA Selectively Modulates Activity of Virulent Amyloid PSMα3 and Host Defense LL-37 via Phase Separation and Aggregation Dynamics

**DOI:** 10.1101/2025.06.17.660072

**Authors:** Bader Rayan, Eilon Barnea, Rinat Indig, Christian F. Pantoja, Jesse Gayk, Yael Lupu-Haber, Alexander Upcher, Amir Argoetti, Jacob Aunstrup Larsen, Alexander K. Buell, Markus Zweckstetter, Meytal Landau

## Abstract

Amyloid-forming peptides are increasingly recognized as dynamic regulators at the host–pathogen interface, yet how environmental factors control their assembly and activity remains poorly understood. Here, RNA acts as a concentration-dependent regulator of two sequence-related α-helical peptides with fundamentally different assembly behaviors: the cross-α amyloid-forming *Staphylococcus aureus* virulence factor PSMα3 and the non-amyloidogenic human host-defense peptide LL-37. RNA drives PSMα3 through distinct assembly states, from liquid-like condensates to fibrillar polymorphs, while preserving cytotoxic and antimicrobial activity over time. In contrast, RNA attenuates LL-37 cytotoxicity toward host cells while maintaining antibacterial activity, consistent with a host-protective immunomodulatory effect. Together with the opposing effects of EGCG, which redirects both peptides into amorphous assemblies, these findings support a mechanistic model in which biological activity is governed by supramolecular architecture, assembly trajectory, and dynamics rather than by monomer abundance or mature fibrils alone. More broadly, our findings identify RNA as an environmental regulator of α-helical peptide assemblies and establish assembly-state control as a tunable determinant of virulence and host defense.

## Introduction

Bacterial amyloids are proteins capable of forming ordered fibrils, playing a crucial role in biofilm stability, surface adherence, cell toxicity and immune responses (Barnhart and Chapman, 2006; Howard et al., 2023; Peña-Díaz et al., 2024; Salinas et al., 2020; Sleutel et al., 2023; Van Gerven et al., 2018). These amyloids are vital for the survival and virulence of many bacteria, enabling them to colonize and persist in diverse environments (Blanco et al., 2012; Chapman et al., 2002). Some examples of bacterial amyloids include the Csg and Fap proteins from *Enterobacteria* and *Pseudomonas* species (Barnhart and Chapman, 2006; Chapman et al., 2002; Golan et al., 2025; Jiang et al., 2025; Peña-Díaz et al., 2024; Perov et al., 2019), respectively, which play a role in biofilm stability. Staphylococcal phenol-soluble modulins (PSMs) are short peptides that predominantly adopt α-helical conformations in solution (Towle et al., 2016) but are capable of forming amyloid fibrils (Hansen et al., 2024; Kreutzberger et al., 2022; Schwartz et al., 2012, 2014; Schwartz and Boles, 2013; Tayeb-Fligelman et al., 2017, 2020; X. Wang et al., 2023; Zhou et al., 2021), and play multiple roles central to pathogenesis and immune evasion in *Staphylococcus aureus* (Arad et al., 2023; Chatterjee et al., 2013; Cheung et al., 2014a; Grando et al., 2022; Hu et al., 2022; Joo and Otto, 2012; Kretschmer et al., 2010; Le et al., 2014; Marinelli et al., 2016; Mehlin et al., 1999; Peschel and Otto, 2013; Queck et al., 2009; Rayan et al., 2023; Zaman and Andreasen, 2020). In this context, defining PSMs as “functional amyloids” implies that amyloid formation can serve regulatory purposes, including controlled storage, stabilization, or spatial organization of bioactive molecules, as described for peptide hormones (Seuring et al., 2020) and RNA-binding proteins (Banerjee et al., 2017; Morelli et al., 2024). Accordingly, amyloid assembly may function as a reversible structural state that modulates the availability and timing of virulence-associated activities.

Among bacterial amyloids, PSMα3 is notable for its strong cytotoxicity toward a broad range of microbial and human cell types (Cheung et al., 2014a, 2014b; Laabei et al., 2014; Peschel and Otto, 2013; Wang et al., 2007) and for adopting a unique cross-α amyloid architecture, in which α-helices stack perpendicular to the fibril axis in paired sheets (Kreutzberger et al., 2022; Tayeb-Fligelman et al., 2020, 2017). Within the PSMα family, distinct amyloid polymorphs appear to encode different biological functions (Rayan et al., 2023): the metastable and reversible cross-α assemblies of PSMα3 may enable dynamic switching between inactive and active states, whereas the more stable and irreversible cross-β fibrils formed by the homologous PSMα1 peptide (Hansen et al., 2024; Salinas et al., 2018) may contribute to robust biofilm architecture. Two-dimensional infrared spectroscopy has further revealed that PSMα3 forms a heterogeneous ensemble of interconverting cross-α and cross-β assemblies, rather than a single defined structure (Cracchiolo et al., 2022)

The cytotoxic mechanism of PSMα3 remains debated, with studies attributing activity to soluble monomers (Yao et al., 2019; Zheng et al., 2018), or implicating the fibrillation process as required for full cytotoxic activity (Malishev et al., 2018; Tayeb-Fligelman et al., 2017). Notably, fibril-forming antimicrobial peptides show a correlation between α-helical conformations in the solid state and cytotoxicity (Bücker et al., 2022; Landau et al., 2026; Ragonis-Bachar et al., 2022; Salinas et al., 2021; Strati et al., 2025), paralleling human amyloids, where α-helical oligomeric intermediates are widely considered the primary toxic species (Ghosh et al., 2015; Ramamoorthy and Lim, 2013; Tempra et al., 2021). Together, these observations support a model in which toxicity arises from transient assembly intermediates along the fibrillation pathway rather than from monomers or mature fibrils alone (Cali et al., 2026).

PSMα3 shares self-assembly of α-helices with LL-37, a human host-defense peptide (Porcelli et al., 2008; Sancho-Vaello et al., 2020; Zeth and Sancho-Vaello, 2021), even though LL-37 is not an amyloid and does not form cross-α or cross-β fibrils. Moreover, the LL-37 antimicrobial active core (residues 17–29) shows sequence similarity to PSMα3 and forms non-amyloid fibrils composed of densely packed cationic amphipathic α-helices (Engelberg and Landau, 2020). The sequence similarity and α-helices self-assembly behavior supported the hypothesis that PSMα3 might recapitulate functions of LL-37, and indeed, they share the activation of the signal inhibitory receptor on leukocytes-1 (SIRL-1), a negative regulator of myeloid cell function and dampens antimicrobial responses (Rumpret et al., 2021). Such common functions might be related to molecular mimicry and the ability of bacterial peptides to hijack immunomodulation pathways (Golan et al., 2022). Indeed, PSMαs interact with the formyl peptide receptor 2 (FPR2) on immune cells, initiating immune responses such as chemotaxis, phagocytosis, and the secretion of pro-inflammatory cytokines. These reactions are critical for recruiting and activating immune cells, thereby amplifying the inflammatory response (Bröker et al., 2016; Chu et al., 2018; Kretschmer et al., 2012, 2010; Rautenberg et al., 2011; Schreiner et al., 2013; Syed et al., 2015).

LL-37 has been shown to bind RNA released from dying cells, forming stable complexes that protect the RNA from enzymatic degradation and promote its uptake by dendritic cells (Ganguly et al., 2009). Once internalized, these RNA–LL-37 complexes activate Toll-like receptors (TLRs), triggering cytokine production and dendritic cell maturation (Ganguly et al., 2009; Lee et al., 2017). Beyond LL-37, other human AMPs, such as β-defensins, also form higher-order assemblies that organize nucleic acids into nanostructures with defined periodicity. These nanocomplexes potently stimulate TLRs, highlighting that innate immune activation is influenced not only by ligand identity, but also by the nanoscale geometry and spatial organization of the complexes (Lee et al., 2019; Schmidt et al., 2015; Tewary et al., 2013).

Interestingly, nucleic acid interactions also play a key role in the immune activation triggered by bacterial amyloids. For example, PSMs co-assemble with extracellular DNA (eDNA), and CpG-rich DNA specifically promotes PSMα3 fibrillation, forming co-aggregates that colocalize with DNA in biofilms. These complexes activate TLR pathways and induce anti-dsDNA autoantibody responses in mice (Grando et al., 2022). Similar mechanisms are seen in *E. coli* and *Salmonella*, where curli fibers bind eDNA, enhance fibrillation, and engage TLR signaling, also promoting autoimmunity (Gallo et al., 2015; Tursi et al., 2017). These findings suggest that amyloid–DNA complexes in bacterial biofilms are not only structural components but also potent immunomodulators with potential links to autoimmunity.

Given their shared cationic, amphipathic α-helical character, but distinct amyloidogenic properties, we sought to examine whether RNA differentially influences the assembly landscapes and bioactivity of PSMα3 and LL-37. RNA is increasingly recognized as a key regulator of protein phase behavior, capable of promoting liquid–liquid phase separation (LLPS), altering aggregation pathways, and reshaping supramolecular organization. LLPS is a fundamental mechanism of cellular organization, in which biomolecules demix into dynamic, membraneless compartments that regulate processes such as RNA metabolism, stress responses, and signal transduction (Alberti et al., 2019; Hyman et al., 2014; Wang et al., 2021; Xu et al., 2023). Whether bacterial virulence-associated peptides are similarly governed by RNA-mediated phase behavior remains largely unexplored.

To address this, we investigated how defined RNA species modulate the structural transitions, material properties, and bioactivity of PSMα3 and LL-37. In parallel, we examined the impact of epigallocatechin gallate (EGCG), a small-molecule assembly modulator widely studied as an amyloid inhibitor (Andrich and Bieschke, 2015; Fernandes et al., 2021; Indig and Landau, 2023; Marinelli et al., 2016), to compare RNA-driven regulation with pharmacological redirection of peptide assembly. Here we show that RNA acts as a concentration-dependent modulator of PSMα3 and LL-37 assembly, inducing LLPS at low concentrations and distinct aggregated states at higher concentrations, with divergent functional consequences that reveal assembly architecture as a tunable determinant of peptide bioactivity.

## Results

### PSMα3 interacts with RNA in vitro

The interactions between PSMα3 and RNA were assessed using an Electrophoretic Mobility Shift Assay (EMSA), comparing the binding affinity of PSMα3 to single-stranded PolyA RNA and double-stranded Poly (AU) RNA. These RNAs were selected as simplified, well-defined model RNAs to probe general peptide–RNA interactions in an unbiased manner, as no prior information was available regarding whether such interactions occur, or which specific RNA species might be involved. The RNA molecules were labeled with the IR800CW fluorescent dye (as detailed in the Methods section). Freshly dissolved PSMα3, at varying concentrations, was incubated with each RNA type at a constant concentration for 30 minutes at 37°C. The EMSA results revealed that PSMα3 exhibits a stronger binding affinity to double-stranded Poly (AU) RNA, as evidenced by significant shifts at lower peptide concentrations compared to single-stranded Poly (A) (Fig. 1).

**Figure 1.**
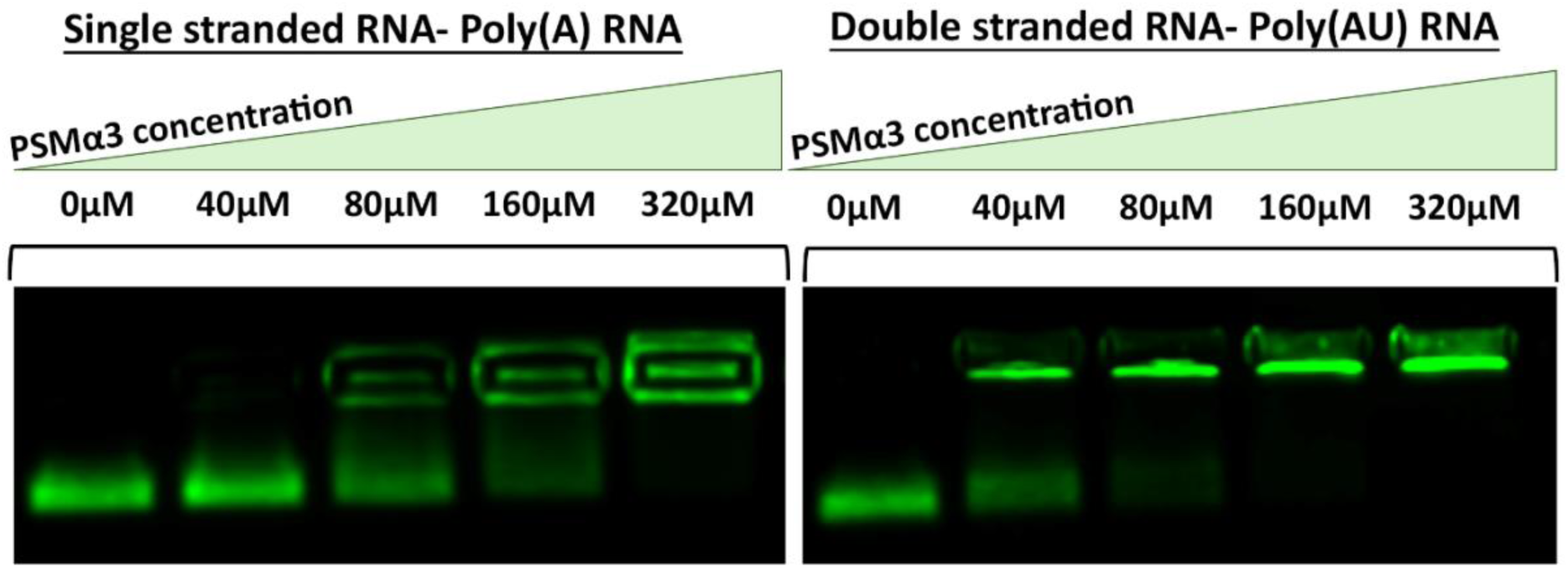
The binding and effect of single- and double-stranded RNA on PSMα3. EMSA assay illustrating the interaction between PSMα3 and RNA. The assay compares the effects of increasing concentrations of PSMα3 (0 µM, 40 µM, 80 µM, 160 µM, and 320 µM), shown in a gradient from left to right, on the mobility shift of single-stranded Poly (A) RNA (left panel) and double-stranded Poly (AU) RNA (right panel) at ∼400ng/µL.

Quantification of free and shifted RNA band intensities revealed a progressive increase in the fraction of RNA bound as a function of peptide concentration for both RNA molecules (Fig. S1). Poly (AU) RNA reached saturation at lower PSMα3 concentrations than Poly (A), consistent with stronger apparent binding. Fitting the binding curves to a Hill model yielded an apparent dissociation constant (Kd) of 61.8 ± 1.6 µM for Poly (AU) and 94.5 ± 5.4 µM for Poly (A) (Fig. S1). Hill coefficients greater than 1 were obtained for both RNAs, consistent with cooperative binding behavior. Based on these Kd differences, subsequent experiments focused on PSMα3 behavior in the presence of double-stranded Poly (AU) RNA (hereafter referred to as RNA).

### Non-monotonic turbidity response of PSMα3 to RNA

In order to assess the concentration-dependent effect of RNA on PSMα3 we used turbidity assays. The results showed that a 30 min co-incubation of 100 µM PSMα3 with varying concentrations of RNA displayed an increased turbidity up to 50 ng/µl, while higher RNA concentrations displayed a decrease in turbidity (Fig. S2). The decrease in turbidity above 50 ng/µL can be a result of either the formation of smaller particles or aggregation into larger clusters, which can settle out of suspension, removing them from the light-scattering medium and reducing turbidity. Alternatively, this non-monotonic turbidity behavior is conceptually similar to charge-driven re-entrant transitions described in other protein–RNA phase-separating systems, such as Ddx4(Nott et al., 2015), however, the molecular context and assembly architecture of PSMα3 are fundamentally distinct. According to the turbidity results, we hypothesized that 50 ng/µl can be a critical concentration between soluble species, which might be able to phase separate, and the formation of aggregates.

### RNA concentration regulates formation and liquid-to-solid transition of PSMα3 condensates

To better understand the nature of the PSMα3-RNA interaction, we visualized their mixtures via fluorescence microscopy, using PSMα3 labeled with Fluorescein isothiocyanate (FITC) at the C-terminus (PSMα3-FITC), which is known to form fibrils (Tayeb-Fligelman et al., 2020). All fluorescence microscopy experiments used a mixture of 20% FITC-labeled and 80% unlabeled peptide to minimize potential effects of the fluorophore on assembly behavior. 100 µM PSMα3-FITC was incubated with either 50 ng/µl or 400 ng/µl RNA, labeled using propidium iodide (PI), in a near-physiological buffer of 50 mM HEPES (pH 7.4) containing 150 mM sodium chloride (NaCl).

Upon addition of 50 ng/µL RNA, PSMα3 formed liquid–liquid phase-separated droplets that clearly colocalized with RNA (Fig. 2A). We further observed encapsulation of RNA within PSMα3–FITC–coated condensates (Fig. S3), indicating co-assembly rather than simple surface association. The liquid-like nature of these droplets was confirmed by fluorescence recovery after photobleaching (FRAP), which showed rapid signal recovery shortly after bleaching (Fig. 2B). Quantitative analysis revealed a half-time of recovery of 3.1 seconds and a mobile fraction of 68% (immobile fraction 32%) (Fig. S4), consistent with dynamic molecular exchange. Across multiple samples, we consistently observed coexistence of small droplets and larger aggregates. The experimental timescales examined do not allow us to reliably determine whether diffusion-driven coalescence kinetics would support classical droplet ripening dynamics (Amico et al., 2024; Ray and Buell, 2024). Notably, after 2 hours of incubation, fluorescence recovery was no longer detected (Fig. 2C), indicating that the condensates undergo time-dependent aging and transition toward a more solid-like or aggregated state.

**Figure 2.**
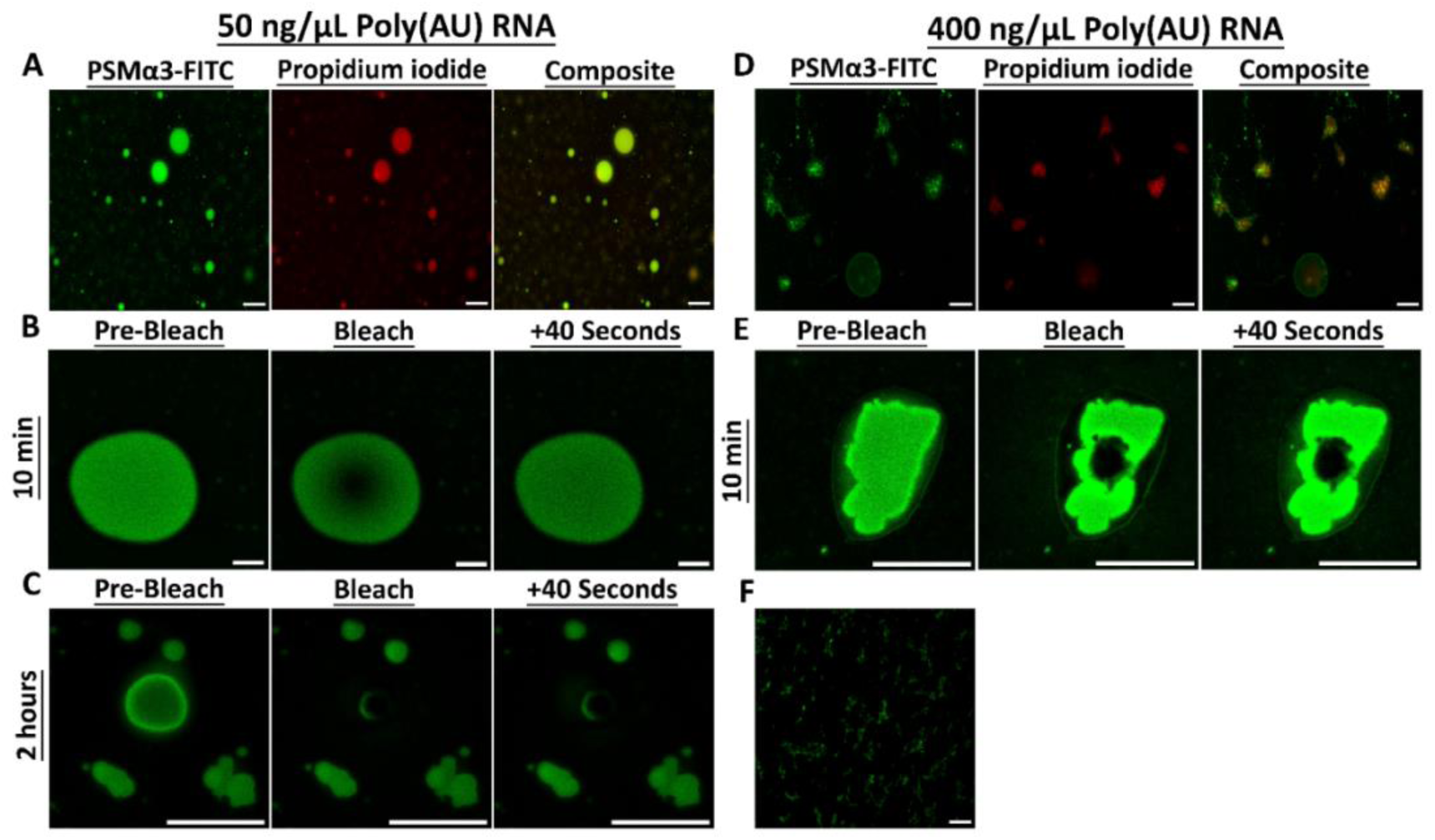
Colocalization, droplet formation, and texture of PSMα3 mixed with varying Poly (AU) RNA concentrations. Widefield fluorescence microscopy images of 100 µM of 20% FITC-PSMα3 (green) and 80% unlabeled PSMα3 incubated with Poly (AU) RNA (red) at 50 ng/µL (**A**) or 400 ng/µL (**D**), shown as individual fluorescence channels and merged images. (**B-C**) FRAP analysis of PSMα3-FITC in the presence of 50 ng/µL Poly (AU) RNA after 10 min (**B**) or 2 h (**C**) of co-incubation. (**E**) FRAP analysis of PSMα3-FITC in the presence of 400 ng/µL Poly (AU) RNA after 10 min of co-incubation. For panels B, C, and E, images were acquired before photobleaching, immediately after photobleaching, and 40 s post-photobleaching. (**F**) 100 µM of 20% FITC-PSMα3 (green) and 80% unlabeled PSMα3 in the absence of RNA. All scale bars represent 20 µm.

In contrast to the effect of 50 ng/µl RNA, the addition of a higher RNA concentration of 400 ng/µl to PSMα3 resulted in immediate aggregation, with no observed phase of droplet formation (Fig. 2D). FRAP analysis indicated no recovery of signal after bleaching, likely due to the formation of solid structures rather than droplets with liquid-like properties (Fig. 2E). In the absence of RNA, PSMα3 did not form condensates and instead appeared as dispersed or irregular assemblies (Fig. 2F), establishing that RNA is required to induce the droplet state observed under these conditions.

Colocalization between PSMα3-FITC and RNA was quantified using Pearson’s correlation coefficient (above the Costes threshold) and Manders’ overlap coefficients (tM1 and tM2) (Fig. S5). All metrics indicate substantially stronger spatial correlation and signal overlap at 50 ng/µL RNA compared with 400 ng/µL RNA. At low RNA concentration, Pearson’s correlation is positive (R ≈ 0.53), accompanied by high Manders’ overlap values (tM1 ≈ 0.68; tM2 ≈ 0.63). This is consistent with coordinated association of PSMα3 and RNA within liquid-like droplets. In contrast, at 400 ng/µL RNA, Pearson’s correlation becomes negative (R ≈ −0.24) and Manders’ coefficients are markedly reduced. This indicates a transition from co-localized droplet assemblies to spatially segregated aggregated structures in which RNA becomes partially excluded from the peptide-rich structures.

### RNA concentration-driven changes in PSMα3 fibrillar morphology and aggregation

The TEM micrographs showed that in the absence of RNA, 100 µM PSMα3 formed nanotube-like fibrils after 2 hours, growing wider or with some twist after 24-hours co-incubation (Fig. 3A). Incubating PSMα3 with a low concentration of 10 ng/µL RNA displayed accelerated fibril formation, with more strongly twisted, wide, sheet-like fibrils observed after both 2 hours and 24 hours (Fig. 3A). With the addition of 50 ng/µL RNA, PSMα3 fibrils have a similar twisted morphology after 2 hours, while at a longer incubation time of 24 hours, we observed a significant morphological shift into more thin amorphous aggregates (Fig. 3A). At the higher concentration of 400 ng/µL RNA, PSMα3 formed dense, thin fibrils after 2 hours, but with a possible fragmentation into smaller species or rearrangement into amorphous aggregates after 24 hours of co-incubation. This indicates that RNA concentration and time of co-incubation affect the density and morphology of PSMα3 fibrils. This corresponds to the differences observed by light microscopy of co-aggregates morphology of LLPS droplets vs solid aggregates (Fig. 2).

**Figure 3.**
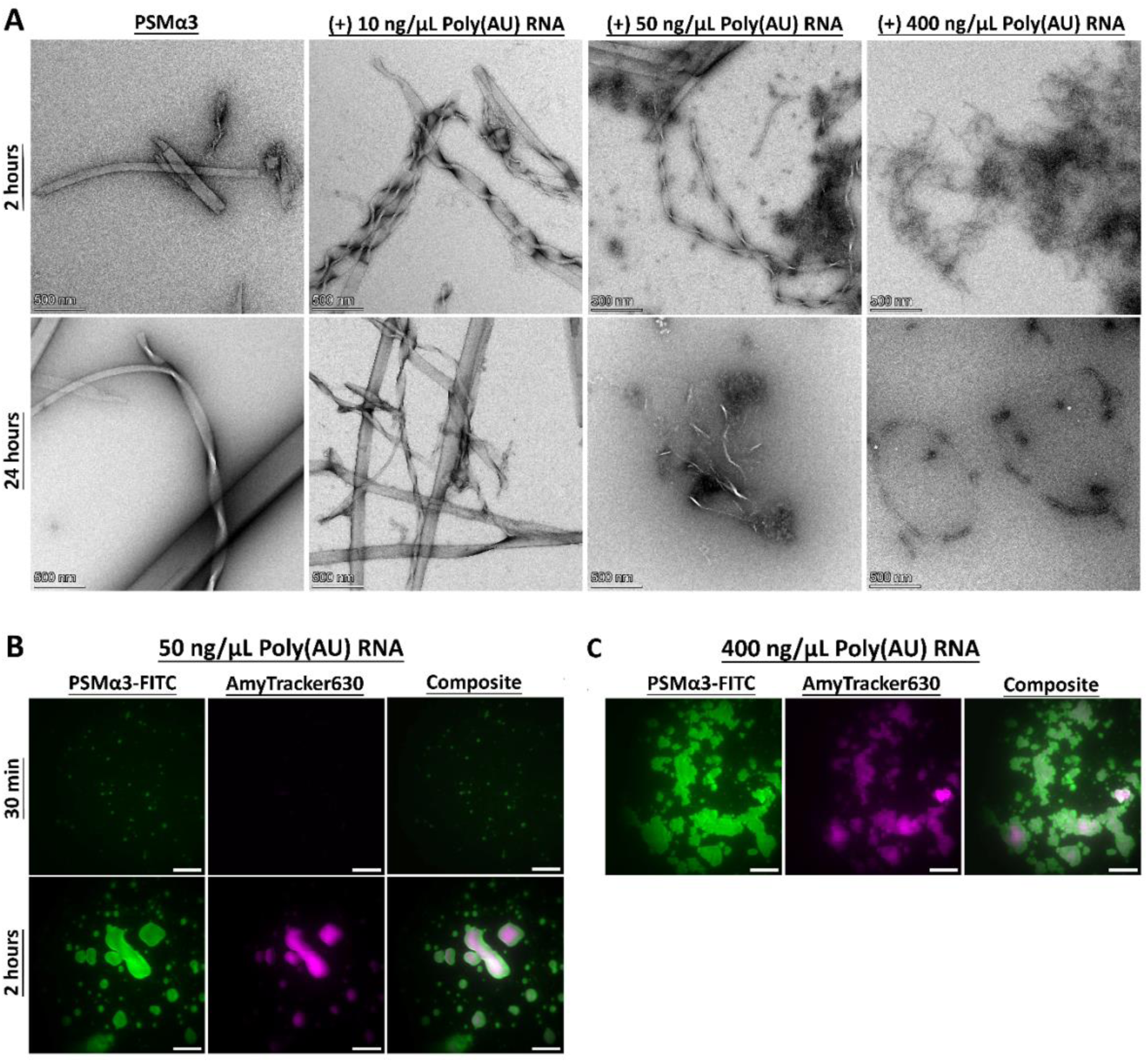
TEM and TIRF visualization of PSMα3 aggregation and morphology with different Poly (AU) RNA concentrations and incubation times. **(A)** TEM micrographs of 100 µM PSMα3 incubated with or without Poly (AU) RNA at varying concentrations of 10 ng/µL, 50 ng/µL, and 400 ng/µL for 2 hours (top row) and 24 hours (bottom row). Scale bars represent 500 nm. **(B)** TIRF microscopy images showing 100 µM of 20% FITC-PSMα3 (green) and 80% unlabeled PSMα3 co-incubated with 50 ng/µL Poly (AU) RNA and the amyloid indicator AT630 (magenta) for 30 minutes and 2 hours. Scale bars represent 20 μm. **(C)** TIRF microscopy images of 100 µM of 20% FITC-PSMα3 (green) and 80% unlabeled PSMα3 co-incubated with 400 ng/µL Poly (AU) RNA and AT630 (magenta) for 30 min. Scale bars represent 20 µm.

These observations were further supported by TIRF microscopy using the amyloid-specific dye AmyTracker630 (AT630). With the addition of 50 ng/µL RNA to PSMα3-FITC, a strong AT630 fluorescence signal was detected only after 2 hours of co-incubation, but not after 30 minutes (Fig. 3B, quantified in Fig. S6). This suggests a time-dependent transition into amyloid-like species. Conversely, at 400 ng/µL RNA, a significant AT630 fluorescence was observed already after 30 minutes of co-incubation, consistent with a rapid formation of amyloid fibrils (Figs. 3C&S6). These findings highlight a concentration- and time-dependent modulation of PSMα3 phase separation and structural transitions, where RNA promotes LLPS at lower concentrations and drives rapid amyloid formation and unique fibrillar morphologies at higher concentrations. Future work will be required to quantitatively define the phase boundaries and delineate the dominant mechanisms, such as sedimentation, dissolution, or coarsening/aging, across intermediate RNA concentrations.

### RNA enhances α-helical structure of PSMα3

Solid-state circular dichroism (ssCD) spectroscopy reveals that PSMα3 lacks a defined secondary structure both immediately after preparation and following 2 hours of incubation (Fig. S7). However, an α-helical signature of PSMα3 is markedly enhanced in the presence of RNA compared to peptide alone, as evidenced by increased signal intensity, deeper minima, and more pronounced spectral features characteristic of α-helical structure. This enhancement is more pronounced at 400 ng/µL RNA than at 50 ng/µL, particularly after 2 hours of co-incubation, indicating that RNA concentration influences the stabilization of α-helical assemblies. This supports the notion that RNA not only accelerates aggregation but also promotes or stabilizes the α-helical fibrillar architecture, potentially consistent with cross-α amyloid structures. This RNA-stabilized α-helical fibrillar architecture provides a structural basis for understanding why RNA preserves PSMα3 bioactivity over extended incubation times, as shown below.

### RNA modulates the antibacterial activity of PSMα3

The antimicrobial activity of PSMα3 against *Escherichia coli* was evaluated under varying RNA concentrations and incubation times using the PrestoBlue Cell Viability Assay (Fig. 4A). Freshly dissolved 10 or 20 µM PSMα3 exhibited potent antibacterial activity, completely abolishing bacterial viability. After 2 hours of incubation, PSMα3 retained its full activity, comparable to its freshly prepared state. The presence of RNA at 50 ng/µL and 400 ng/µL had no impact on its antimicrobial function within this timeframe (Fig. 4A).

**Figure 4.**
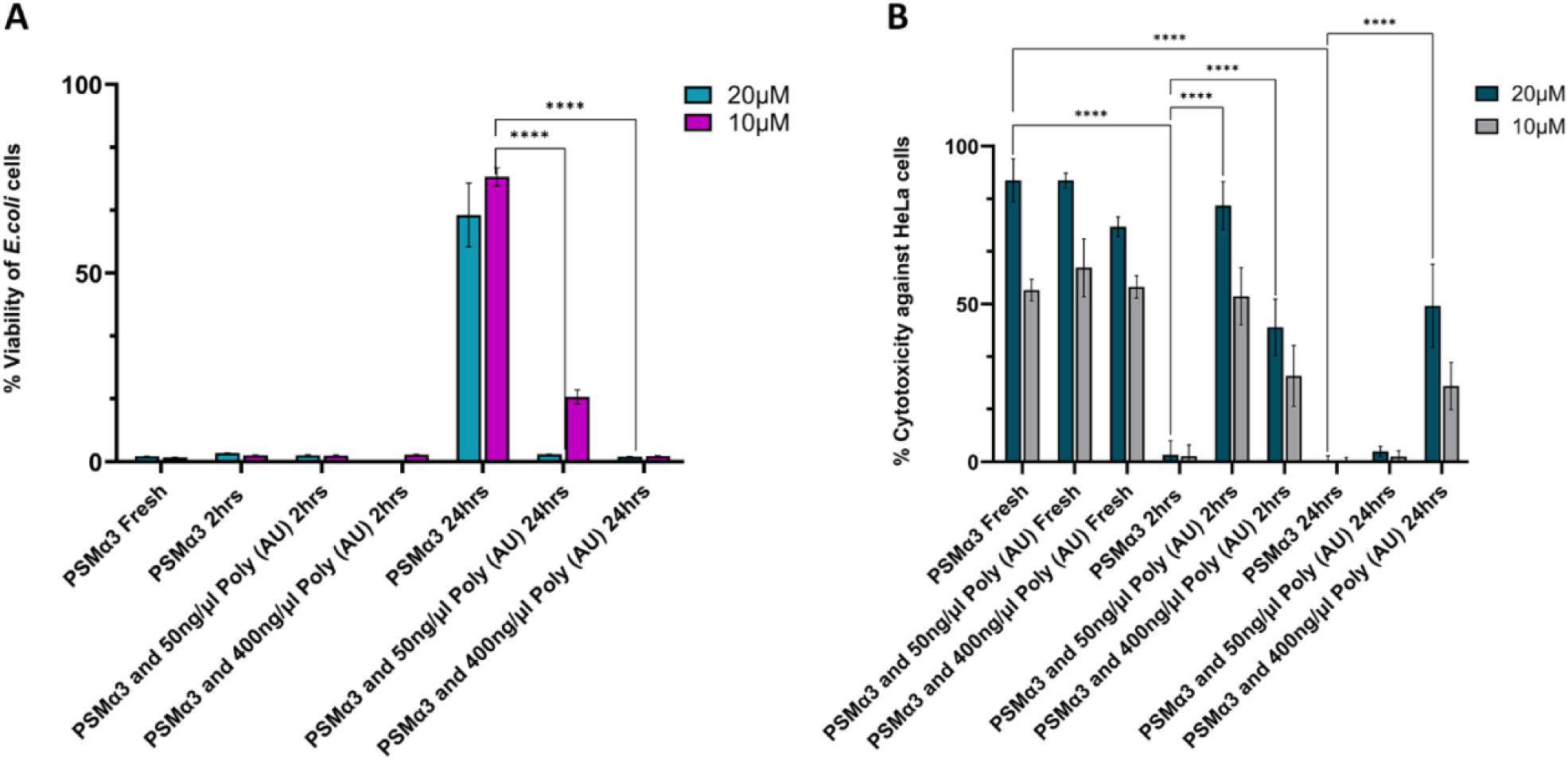
Impact of Poly (AU) RNA on PSMα3 cytotoxicity and antibacterial activity. Antimicrobial activity of PSMα3 against *E. coli* using the PrestoBlue cell viability assay (**A**) and its cytotoxicity against HeLa cells using the LDH colorimetric assay (**B**) were assessed with and without Poly (AU) RNA at varying concentrations. The experiments were performed in at least three replicates and repeated across three independent days to ensure result reliability. Cytotoxicity and bacterial cell viability percentages were calculated as the mean of all replicates, with error bars representing the standard deviation. Statistical significance was determined using one-way ANOVA for normally distributed data in GraphPad Prism (version 11). Significance levels are indicated as follows: *p<0.05, **p<0.01, ***p<0.001, ****p<0.0001.

In contrast, after 24 hours of incubation, PSMα3’s antibacterial activity was significantly reduced (Fig. 4A), suggesting a decrease in its effective concentration or changes in its morphology. Notably, the addition of RNA at 50 ng/µL or 400 ng/µL prevented this loss of activity, indicating that RNA plays a stabilizing role in toxic species or their reservoir. These findings suggest that RNA influences PSMα3 aggregation dynamics and morphology, thereby modulating its long-term antimicrobial effectiveness.

### RNA modulates PSMα3 cytotoxicity against human HeLa cells

The effect of RNA on the cytotoxicity of PSMα3 against human HeLa cells was evaluated by measuring lactate dehydrogenase (LDH) release, an indicator of cell membrane damage (Fig. 4B). Freshly dissolved PSMα3 exhibited substantial cytotoxicity, causing approximately 80% cell death at 20 µM and 50% at 10 µM. The presence of RNA did not significantly alter the toxicity of freshly dissolved PSMα3 (Fig. 4B), similar to the antibacterial activity. However, following 2 hours of incubation, PSMα3 cytotoxicity was significantly reduced, likely due to aggregation-associated loss of membrane-active species. This reduction occurred earlier than the loss of antibacterial activity, which was only observed after 24 hours of incubation (Fig. 4A).

Notably, co-incubation with RNA helped maintain PSMα3 cytotoxicity in a concentration-dependent manner. With 50 ng/µL RNA, the 2-hour incubated PSMα3 retained cytotoxicity comparable to its freshly dissolved form, suggesting that RNA prevents activity loss due to incubation. However, after 24 hours, 50 ng/µL RNA was insufficient to preserve cytotoxicity. In contrast, at 400 ng/µL RNA, partial cytotoxicity was maintained for both 2-hour and 24-hour incubated PSMα3, compared to the freshly dissolved sample.

Overall, these findings suggest that RNA prevents the incubation-induced loss of both antibacterial activity and cytotoxicity (Fig. 4). This effect appears to be concentration- and co-incubation time-dependent, likely linked to RNA-induced morphological variations of PSMα3 species (Figs. 2-3).

### PSMα3 accumulates in nucleolar RNA-rich compartments of HeLa cells

The localization of PSMα3 in HeLa cells was examined following the addition of 20µM PSMα3 immediately before imaging. PI was added to the cell medium as a marker for cell death, as it selectively penetrates dead cells and binds to their nucleic acids. Confocal microscopy images revealed that PSMα3 induces significant toxicity as indicated by membrane damage and blebbing, intracellular PSMα3 aggregate formation, and positive PI staining (Fig. 5 and Movie S1). PSMα3 also enters the nuclei, with a notable concentration in the nucleolus, indicated by the foci of green fluorescence (marked by white arrows in Fig. 5). The composite images demonstrated clear colocalization of PSMα3 with nucleic acids in the nucleolus, as seen by the overlap of the green PSMα3-FITC signal and the red fluorescence from PI-stained nucleic acids.

**Figure 5.**
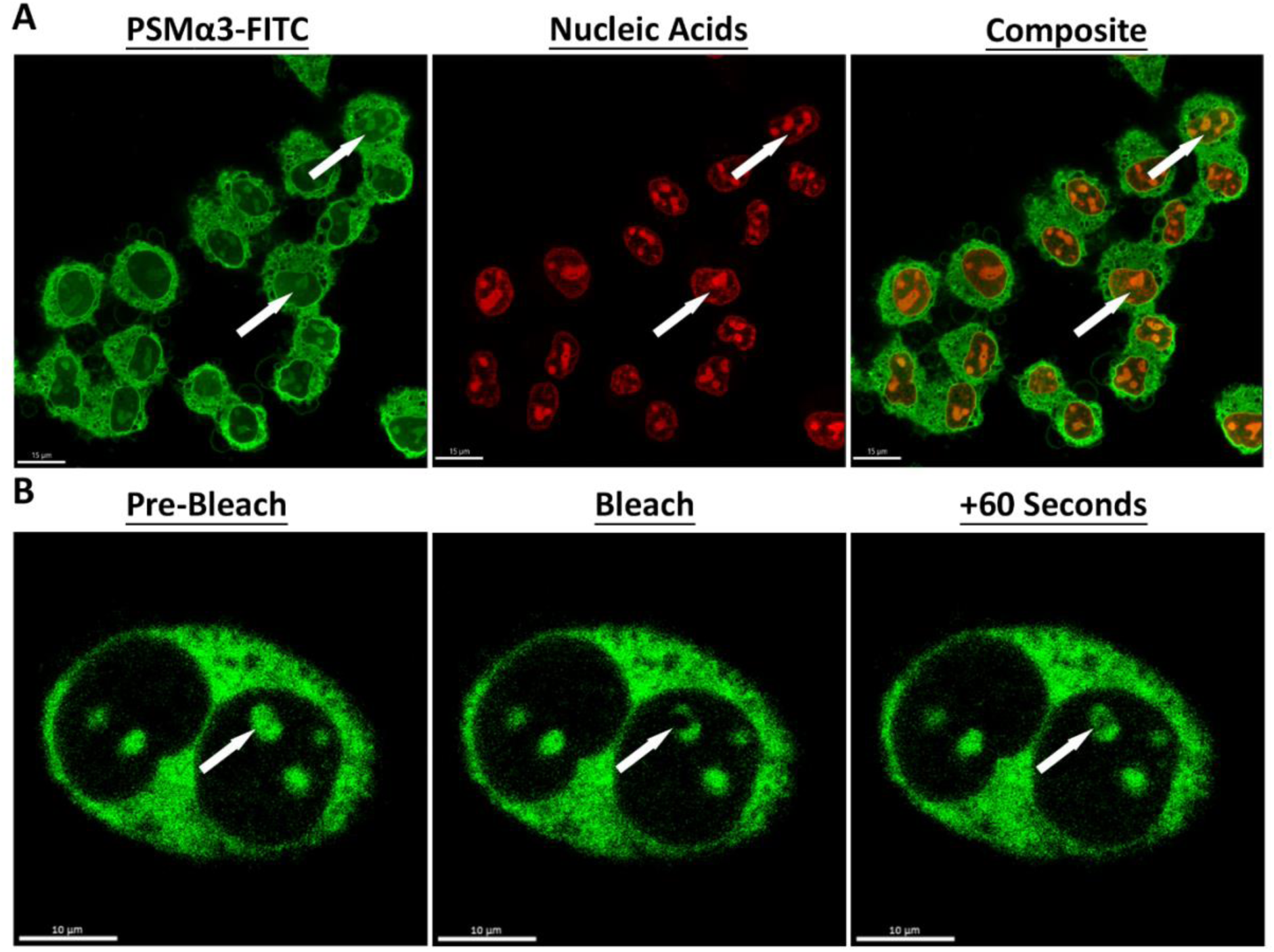
Colocalization of PSMα3 with nucleic acids in HeLa cells. **(A)** Confocal microscopy images showing the localization and colocalization of 20 µM of 20% FITC-PSMα3 (green) and 80% unlabeled PSMα3 and nucleic acids stained with PI (red) inside the nucleolus of HeLa cells (indicated in arrows). The left panel illustrates the distribution of PSMα3 within the cell. The middle panel shows the nucleic acids stained with PI. The right panel is a composite image that demonstrates the colocalization of PSMα3 with nucleic acids. Scale bars represent 15 µm. **(B)** FRAP analysis of 20 µM of 20% FITC-PSMα3 (green) and 80% unlabeled PSMα3 inside the nucleolus (indicated by the arrow) of HeLa cells, showing fluorescence recovery after 60 seconds. Scale bars represent 10 µm.

FRAP analysis of 20 µM PSMα3 within the nucleolus of HeLa cells revealed measurable but constrained mobility (Fig. 5B). Following photobleaching of a defined nucleolar region (indicated by the white arrow), fluorescence gradually recovered over the 60 s acquisition window, indicating dynamic exchange of PSMα3 within this compartment. However, recovery was slow and incomplete during this time frame, and did not reach a clear plateau, precluding reliable determination of recovery half-time or precise mobile and immobile fractions (Fig. S8), and indicating dynamic but constrained exchange within this RNA-rich compartment. This behavior differs from the rapid and near-complete recovery observed for PSMα3–RNA droplets formed in vitro (Figs. 2B&S6), indicating that nucleolar-associated PSMα3 exhibits more restricted dynamics than liquid-like condensates in defined buffer systems.

### EGCG directly binds PSMα3 and inhibits its fibrillation and bioactivity, even in the presence of RNA

Since RNA appears to protect or maintain PSMα3’s toxic functions, potentially by influencing its fibril formation and morphology, we investigated the corresponding and combined effects of an inhibitor of fibril formation. One such inhibitor is EGCG, the most abundant catechin in tea and a known amyloid inhibitor, including PSMs (Marinelli et al., 2016).

The addition of EGCG to PSMα3 at a 1:1 molar ratio did not significantly alter its cytotoxicity against HeLa cells. However, at a fivefold molar excess, EGCG completely abolished PSMα3’s cytotoxic effect (Fig. 6A). To examine the structural changes underlying this effect, we analyzed fibril formation kinetics and morphology in the presence of EGCG. TEM micrographs revealed that a fivefold molar excess of EGCG disrupted fibril formation of 100 µM PSMα3, instead inducing amorphous aggregates (Fig. 6B). Consistently, kinetics assays of fibril formation showed that EGCG inhibited the ThT fluorescence curve otherwise indicating the fibril formation of 100 µM PSMα3 (Fig. S9). These findings suggest that EGCG reduces PSMα3 toxicity by modulating its fibril formation and morphology. The addition of RNA did not counteract the effect of EGCG or restore PSMα3 cytotoxicity (Fig. 6A), highlighting the pronounced effect of EGCG on PSMα3 morphology and properties.

**Figure 6.**
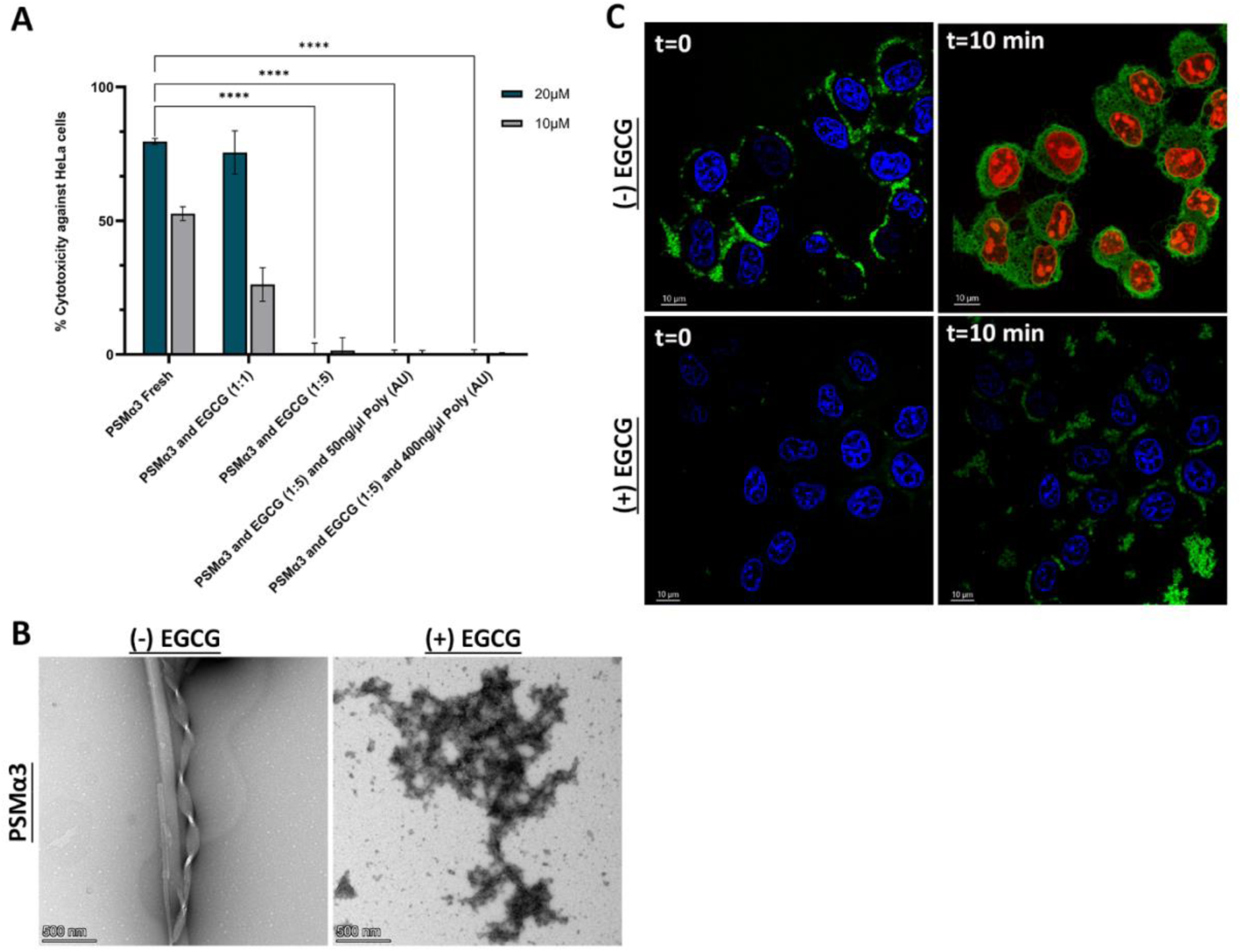
EGCG modulates PSMα3 aggregation and reduces toxicity against HeLa cells. (**A**) Cytotoxicity of PSMα3 against HeLa cells in the presence and absence of EGCG and Poly (AU) RNA at two different concentrations, assessed via LDH assay. The experiment was performed in triplicate and repeated on three separate days for consistency. Cytotoxicity percentages were averaged across all replicates, with error bars representing the standard deviation. Statistical significance: *p < 0.05, **p < 0.01, ***p < 0.001, ****p < 0.0001 (one-way ANOVA, GraphPad Prism v11). (**B**) TEM micrographs of 100 μM PSMα3 incubated for 24 hours, without (left) and with (right) a fivefold molar excess of EGCG. Scale bars: 500 nm. (**C**) Live-cell confocal microscopy of HeLa cells treated with 20 μM of 20% FITC-PSMα3 (green) and 80% unlabeled PSMα3 without (top) and with EGCG (bottom), imaged immediately after preparation (t = 0) and after 10 minutes. Hoechst 33342 (blue) marks the nuclei, while PI staining (red) indicates membrane disruption. Scale bars: 10 μm.

Live-cell confocal microscopy provided further insights into how EGCG binding affects PSMα3 interactions with cells. In accordance with the LDH measurements (Fig. 6A), in the absence of EGCG, 20 µM PSMα3 readily penetrated cells and induced robust PI staining, indicating membrane disruption (Fig. 6C, top panel and Movie S1). In contrast, a fivefold molar excess of EGCG prevented the toxic effects of 20 µM PSMα3, blocking cell penetration and PI staining (Fig. 6C, bottom panel and Movie S2). Instead, PSMα3-FITC formed extracellular aggregates and showed no interaction with cell membranes, as indicated by the absence of PI staining and intact cell membranes.

Furthermore, EGCG at a fivefold molar excess also reduced the antibacterial activity of PSMα3 against *E. coli*, preserving bacterial cell viability (Fig. S10). Super-resolution fluorescence microscopy showed that while 20 µM PSMα3 typically aggregates on the bacterial membrane, causing membrane disruption and PI staining indicative of cell death (Fig. S10), EGCG at a fivefold molar excess prevented membrane aggregation and PI staining, thereby preserving membrane integrity.

To investigate residue-specific interactions between PSMα3 and EGCG, we performed NMR spectroscopy using a 2:1 PSMα3:EGCG molar ratio. The 1D 1H-NMR spectrum revealed distinct chemical shift changes and peak broadening upon EGCG addition, particularly in residues Met1, Glu2, and V4/N21, suggesting specific interactions between these sites and the EGCG molecule (Fig. 7A). Peak broadening was also observed in the aromatic region of EGCG, especially for protons H1 and H2, compared to the reference spectrum of EGCG alone, indicating that the interaction is visible from the EGCG side as well. Notably, slight opalescence was observed in the sample following preparation, potentially reflecting early-stage aggregate formation. Additionally, the presence of multiple cross-peaks between non-sequential residues (i+3 or i+4) in the two-dimensional (2D) ^1^H-^1^H Nuclear Overhauser Effect Spectroscopy (NOESY) spectrum indicates a well-defined structure of PSMα3 in this condition (Fig. 7B). The 2D ^1^H-^1^H Total Correlation Spectroscopy (TOCSY) spectrum displayed connectivity between backbone amide (HN) and alpha protons (Hα) for several assigned residues, while the 2D ^1^H-^1^H NOESY spectrum revealed spatial correlations between nearby residues (i+1).

**Figure 7.**
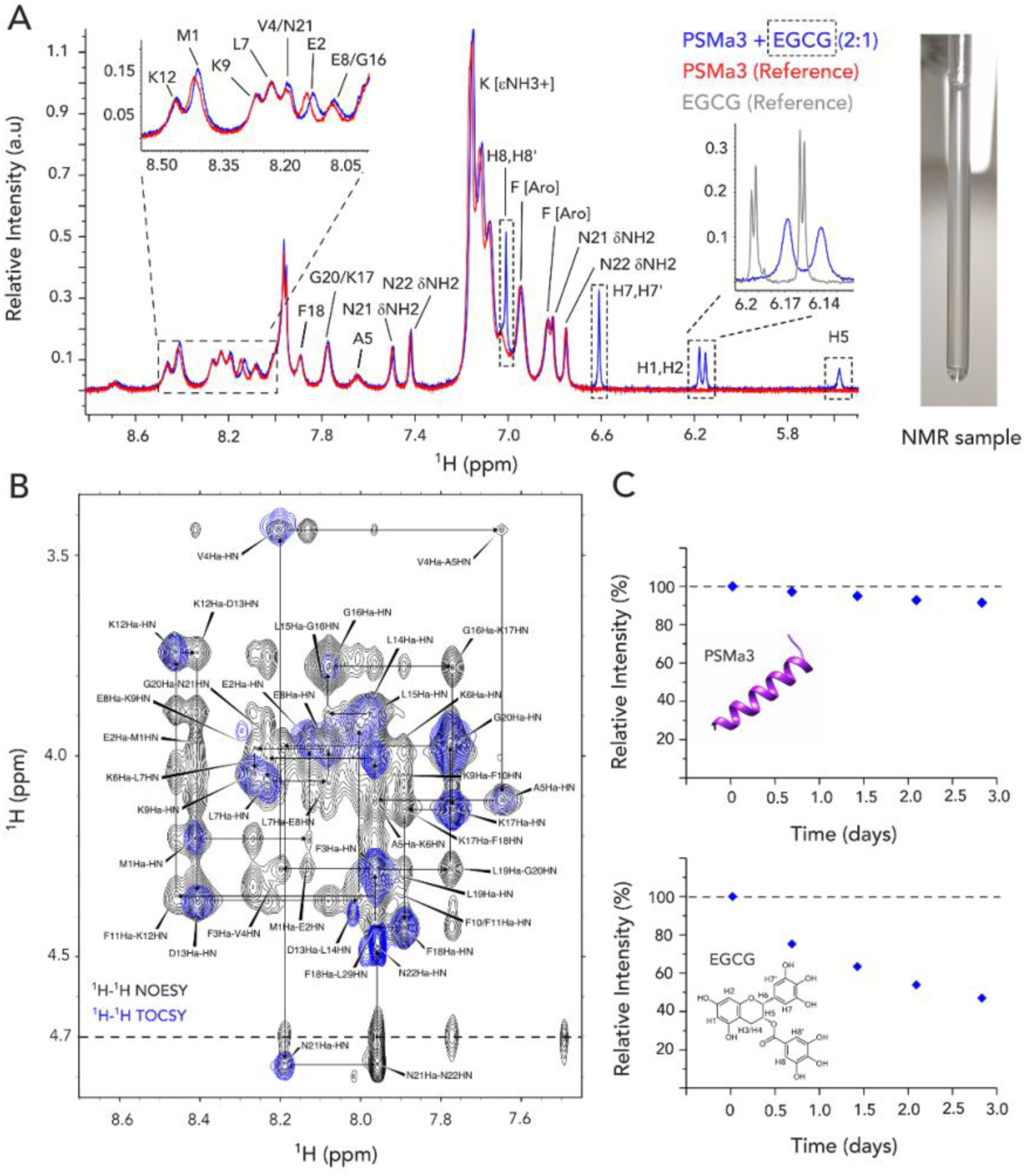
Residue-specific interactions between PSMα3 and EGCG. (**A**) One-dimensional (1D) ^1^H NMR spectra of 1.0 mM PSMα3 alone (red) and in complex with 0.5 mM EGCG (blue), recorded at 35 °C. Specific residues, including Met1, Glu2 and V4/N21, show chemical shift changes suggestive of direct interaction with EGCG (highlighted in the upper left). Dashed boxes mark proton signals corresponding to EGCG. Peak broadening of H1/H2 protons, compared to the EGCG-only reference sample (light grey), indicates interaction from the EGCG side. Slight opalescence observed in the sample suggests potential aggregate formation. (**B**) Two-dimensional (2D) ^1^H–^1^H TOCSY and NOESY spectra of the PSMα3:EGCG complex at a 2:1 ratio, recorded at 35 °C. The cross-peaks in the TOCSY spectrum allow the identification of the spin system and direct connections through the scalar coupling between the proton amide (HN) and the alpha protons (Hα) of the same residue. The cross-peaks in the NOESY spectrum establish a sequential connection between neighboring residues (i+1), allowing spectral assignment. The remaining cross-peaks (i+3 or i+4) in the spectrum support PSMα3 secondary structure in the experimental conditions. (**C**) Temporal stability of the PSMα3:EGCG sample over 3 days.

To assess the temporal stability of the sample, we monitored signal intensities over a 3-day period. PSMα3 signals remained stable, with only a ∼10% decrease in intensity, whereas EGCG signals exhibited substantial degradation, with nearly 50% loss over the same time frame (Fig. 7C). Given the ∼1.5-day duration of the 2D NMR measurements, it is estimated that over 60% of EGCG remained in solution during data acquisition, allowing for reliable observation of its interaction with the peptide. Of note, EGCG’s activity has been shown to depend on its chemical stability and the surrounding conditions. Specifically, at neutral pH, EGCG may undergo oxidation, and its inhibitory effects could be attributed to its degradation products rather than the intact compound itself (Sternke-Hoffmann et al., 2020).

### RNA modulates the human host-defense peptide LL-37 phase behavior and cytotoxicity without compromising antimicrobial activity

LL-37, a human host defense peptide, shares similarities with PSMα3, including its ability to self-assemble into α-helical supramolecular structures and to interact with nucleic acids(Engelberg and Landau, 2020; Ganguly et al., 2009). We therefore examined whether RNA similarly influences LL-37 activity and properties.

Fluorescence microscopy showed that 100 µM LL-37-FITC incubated with 100 ng/µL PI-labeled Poly (AU) RNA formed aggregates, with no evidence of droplet formation or LLPS under these conditions (Fig. S11). Because diverse cellular stresses are known to induce condensation of RNA-binding proteins and promote the formation of stress-associated assemblies (Riback et al., 2017), we next examined the effect of RNA on LL-37 under thermal stress. Following heat shock at 65 °C for 15 minutes, RNA markedly altered LL-37 assembly behavior in a concentration-dependent manner. At 100 ng/µL RNA, LL-37 formed rounded condensates that colocalized with RNA (Fig. 8A). To assess the material properties of these assemblies, we performed FRAP analysis on LL-37–FITC under this condition. Photobleaching resulted in minimal fluorescence recovery over the acquisition window (Fig. S12), indicating that the LL-37–RNA assemblies formed after heat shock are predominantly solid-like rather than liquid-like. At 200 ng/µL RNA, LL-37 assemblies exhibited mixed features, with rounded structures still present but accompanied by increasing aggregation, consistent with a transition toward a more solid state (Fig. 8B). At 400 ng/µL RNA, discrete condensates were no longer observed and LL-37 instead formed extensive, irregular aggregates with strong RNA colocalization (Fig. 8C).

**Figure 8.**
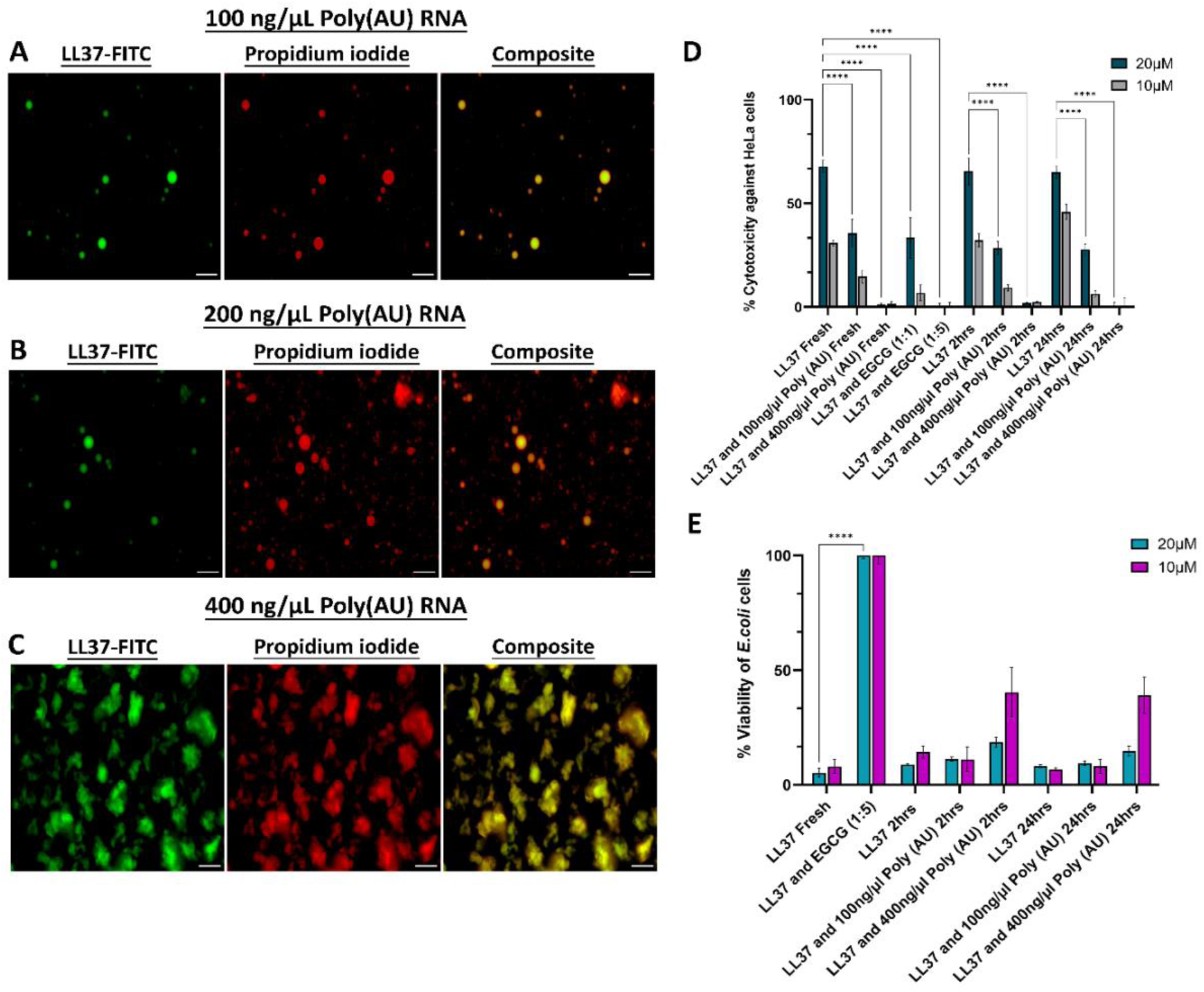
Effect of RNA concentration on LL-37 phase separation, aggregation, and activity at pH 7 after heat shock. Fluorescence microscopy images showing 20% FITC-labeled (green) and 80% unlabeled 100 μM LL-37 in the presence of increasing concentrations: 100 ng/μL (**A**), 200 ng/μL (**B**) and 400 ng/μL (**C**) of Poly (AU) RNA (PI-labeled, red) after heat shock at 65°C for 15 minutes at pH 7. Scale bars represent 20 µm. **(D)** LL-37 cytotoxicity, with and without Poly (AU) RNA and EGCG at varying concentrations, was assessed in HeLa cells using the LDH colorimetric assay. The experiment was performed in triplicate and repeated on three separate days for reproducibility. Cytotoxicity percentages represent the average of all replicates, with error bars indicating the standard deviation. **(E)** The antimicrobial activity of LL-37, with and without Poly (AU) RNA and EGCG at varying concentrations, was evaluated against *E. coli* using the PrestoBlue Cell Viability assay. The experiment was performed in triplicate and repeated on three separate days for reproducibility. Bacterial viability percentages represent the average of all replicates, with error bars indicating the standard deviation. Statistical significance (**D-E**): *p < 0.05, **p < 0.01, ***p < 0.001, ****p < 0.0001 (one-way ANOVA, GraphPad Prism v11).

For comparison, PSMα3 subjected to the same heat-shock protocol exhibited droplet formation only under more permissive conditions (pH 4), but not at physiological pH (7.4) (Figs. S13-S14). Together, these results indicate that, for both LL-37 and PSMα3, increasing RNA concentration favors aggregation over phase separation, and that LL-37 assemblies formed under thermal stress rapidly adopt solid-like properties rather than maintaining liquid-like dynamics.

Freshly dissolved LL-37 exhibited cytotoxicity against HeLa cells, which, in contrast to PSMα3, remained consistent even after incubation for up to 24 hours. Notably, RNA significantly reduced LL-37 cytotoxicity in a concentration-dependent manner, irrespective of the incubation duration (freshly dissolved, 2 hours, or 24 hours) (Fig. 8D). This contrasts with RNA’s rescuing effect on the incubation time-dependent loss of PSMα3 cytotoxicity. The antimicrobial activity of LL-37 against *E. coli* also remained unchanged upon incubation. However, unlike its effect on cytotoxicity, RNA had minimal impact on antibacterial activity, with only high concentrations of 400 ng/µL reducing LL-37 activity (Fig. 8E). Similar to PSMα3, LL-37 cytotoxicity against HeLa cells and its antimicrobial activity were attenuated by EGCG in a concentration-dependent manner (Fig. 8).

TEM micrographs of 100 µM LL-37 revealed a variety of aggregative and fibrous structures, including thin fibrils, although no consistent morphology was observed (Fig. S15A). RNA induced a distinct morphological change, leading to increased aggregation and the formation of mostly amorphous species, but also some thicker, ribbon-like fibrils were observed. These structures resemble those previously observed for a segment of the LL-37 active core (residues 17–29), which is similar to PSMα3 in sequence and its ability to form fibrils of densely packed amphipathic α-helices (Engelberg and Landau, 2020). Heat shock further intensified aggregation, resulting in denser, amorphous condensates and larger fibrillar assemblies, with and without RNA (Fig. S15B). The distinct effects of RNA on the aggregation and morphologies of LL-37 and PSMα3 may underlie the observed differences in their cytotoxic and antibacterial activities.

## Discussion

This study identifies RNA as a context-dependent regulator of α-helical assembly-prone peptides, demonstrating that RNA reshapes their supramolecular landscape and thereby tunes bioactivity rather than merely sequestering peptide species. This is supported by the emergence of distinct RNA-dependent material states (from droplets to aged solids) and RNA-dependent fibrillar morphologies with divergent activity profiles across time and concentration. By comparing the bacterial virulence factor PSMα3, which forms canonical cross-α amyloid assemblies, with the human host-defense peptide LL-37, which forms α-helical fibrils but lacks amyloid-like stacking, we reveal that RNA differentially modulates structurally distinct systems through peptide-specific effects on phase behavior and aggregation dynamics.

### RNA preserves PSMα3 bioactivity by redirecting its assembly pathway

Previous studies have argued that aggregation into amyloids primarily decreases PSM cytotoxicity by depleting the active soluble pool (Yao et al., 2019; Zheng et al., 2018), a view consistent with our observation of time-dependent activity loss upon incubation (Rayan et al., 2023) (Fig. 4), coinciding with the formation of dense, fibrillar assemblies (Figs. 2-3). These mature aggregates likely represent inert end states that sequester membrane-active intermediates (Yao et al., 2019; Zheng et al., 2018). Our data, however, indicate that RNA-dependent modulation of aggregation trajectory can preserve activity independently of simply reducing aggregation, which challenges a purely monomer-centric interpretation. Specifically, RNA reshapes this trajectory in a concentration-dependent manner. At low RNA concentrations, PSMα3 initially retains activity (Fig. 4B) but gradually loses it as liquid-like RNA–PSMα3 condensates mature into more rigid assemblies (Figs. 2-4 & S3), indicating that RNA regulates the lifetime of dynamic intermediates. At higher RNA concentrations, RNA promotes alternative fibrillar polymorphs and is associated with the retention of substantial cytotoxic activity even after prolonged incubation (Fig. 4B). Similarly, RNA preserves antibacterial potency over time (Fig. 4A), consistent with stabilization of assembly states that remain competent for membrane interaction. Thus, RNA does not simply suppress aggregation but potentially modulates the reversibility and supramolecular architecture of α-helical assemblies (Fig. S7), thereby controlling the duration and accessibility of the toxic window.

The temporal differences between cytotoxicity (Fig. 4B) and antibacterial activity (Fig. 4A) may reflect distinct membrane environments, aggregation kinetics, or interactions with bacterial surfaces versus mammalian membranes. Nonetheless, in both contexts, activity correlates with assembly state rather than peptide abundance alone. RNA therefore acts as a structural modulator that influences the lifetime and accessibility of functionally relevant intermediates.

Recent cryo-EM studies show that RNA acts as a structural cofactor in tau fibrils, directing polymorphic folds, stabilizing assemblies via electrostatic interactions, and forming intermediate states linked to pathology; RNase-induced disassembly confirms RNA as an integral architectural component rather than a passive trigger (Abskharon et al., 2026). This parallels our observations for PSMα3, where RNA similarly governs supramolecular architecture and functional outcomes.

### Regulatory modulation versus irreversible inhibition

The contrasting effects of RNA and the amyloid inhibitor EGCG highlight that bioactivity is governed by supramolecular architecture and reversibility rather than aggregation per se. RNA preserves PSMα3 bioactivity by stabilizing dynamic, α-helical assembly states. In contrast, EGCG binds soluble PSMα3 and diverts it into stable amorphous aggregates that represent structural dead ends, abolishing activity (Figs. 6-7). Thus, EGCG should not be viewed as evidence that amyloid fibrils are the toxic species; instead, it abolishes activity by removing or redirecting soluble bioactive species into non-productive amorphous assemblies. If cytotoxic activity were determined solely by free monomer concentration, both RNA and EGCG would be expected to reduce activity through sequestration. Instead, RNA promotes aggregation while preserving activity, whereas EGCG, which also reduces free soluble peptide, abolishes it, demonstrating that the assembly pathway and resulting structural architecture, rather than soluble peptide abundance alone, are determinants of functional outcome. Overall, RNA functions as a regulatory modulator of assembly dynamics, whereas EGCG enforces irreversible pathway diversion. These results underscore that assembly state, not simply peptide presence, determines biological function.

A parallel diversion mechanism is observed with human fibrinogen, an abundant host plasma protein encountered by PSMs at infection and wound sites: it suppresses fibrillation of most PSMs or redirects aggregation into off-pathway species, while uniquely accelerating non-fibrillar, amorphous aggregation of PSMα3 (Najarzadeh et al., 2021). This further illustrates that host environmental factors encountered during infection can reshape PSMα3 assembly trajectories in ways that parallel the RNA- and EGCG-mediated redirections described here.

### PSMα3 cytotoxicity arises from dynamic assembly intermediates

Current evidence supports a model in which PSMα3 cytotoxicity does not arise from a single static species, i.e., neither free monomers nor mature fibrils, but from dynamic assembly processes involving transient intermediates along the fibrillation pathway. PSMα3 assembly is highly sensitive to environmental conditions that shape both its aggregation trajectory and functional output. Lipid bilayers modulate aggregation kinetics in a concentration- and composition-dependent manner without redirecting the dominant fibrillar pathway (Kristoffersen et al., 2024), while hydrodynamic shear comparable to vascular flow accelerates amyloid formation and promotes α-helical fibrils (Zhu et al., 2024). Salt-induced polymorphism further generates structurally distinct amorphous, fibrillar, and oligomeric assemblies from the same peptide sequence, with fibrillar and oligomeric species exhibiting substantially higher cytotoxicity than amorphous aggregates (Xuan et al., 2022), demonstrating that supramolecular architecture, rather than peptide identity or monomer availability, governs activity.

Multiple independent observations converge on membrane-associated intermediates as the primary cytotoxic species. Mutants that retain α-helical monomeric structure yet fail to fibrillate lose toxicity (Tayeb-Fligelman et al., 2020, 2017), while mature fibrils are largely inactive (Yao et al., 2019; Zheng et al., 2018), indicating that neither endpoint state alone accounts for activity. Real-time AFM studies of N-formylated PSMα3, a native modification, further show that protofibrillar intermediates forming at membrane interfaces drive membrane disruption, whereas mature fibrils are less active (Bonnecaze et al., 2024). Consistently, we observe peptide accumulation and aggregation at cellular and bacterial membranes (Figs. 6C and S10), supporting the view that bioactive intermediates are structurally distinct from both soluble monomers and mature fibrils. Together, these findings position the toxic window within transient, membrane-associated assembly states (Cali et al., 2026).

Molecular dynamics simulations further suggest that PSMα3 monomers undergo membrane-induced folding into α-helical conformations upon contact (Wang et al., 2025), implicating the membrane interface as a site of both structural organization and initiation of cytotoxic assemblies. In this framework, membrane-associated fibril growth may destabilize bilayers through lipid recruitment and extraction (Malishev et al., 2018), analogous to mechanisms proposed for membrane-active human amyloids such as islet amyloid polypeptide (Sparr et al., 2004). Together, these findings support a model in which cytotoxicity emerges from dynamic, environmentally regulated assembly intermediates rather than from free monomers or mature fibrils alone.

The RNA-dependent modulation described here represents a specific instance of this broader principle: environmental factors continuously reshape PSMα3 assembly pathways and thereby control the availability, lifetime, and structure of bioactive species. Crucially, RNA does not preserve cytotoxicity by preventing aggregation; it promotes aggregation while altering its trajectory and morphology. RNA-treated samples retain cytotoxicity over extended incubation times, whereas peptide alone loses activity as aggregation proceeds, inconsistent with a model based solely on monomer sequestration. Instead, these findings support a framework in which assembly state, fibril morphology, and aggregation reversibility, rather than free monomer concentration alone, govern cytotoxic function.

### LL-37: non-amyloid α-helical assemblies under RNA control

In contrast to PSMα3, LL-37 does not form canonical amyloid architectures. This structural distinction of amyloidogenic PSMα3 versus non-amyloidogenic LL-37 provides a natural test of whether RNA-mediated phase regulation is specific to amyloid-competent assemblies or represents a broader principle of α-helical peptide regulation. Although LL-37 can assemble into α-helical fibrillar structures (Engelberg and Landau, 2020; Sancho-Vaello et al., 2020), these lack the ordered cross-β or cross-α stacking characteristic of amyloids. Nevertheless, RNA strongly modulates LL-37 behavior. RNA can selectively attenuate cytotoxicity while preserving antimicrobial function, correlating with the formation of amorphous RNA–LL-37 co-assemblies rather than structured fibrils (Figs. 8, S11 & S15). RNA-dependent condensate-like assemblies are observed only under thermal stress, and these rapidly adopt solid-like properties (Figs. 8&S12). For LL-37, RNA therefore appears to serve a protective role, minimizing collateral host damage while maintaining immune defense. This suggests that RNA-mediated modulation is a broader principle of α-helical peptide regulation, yet with a functional divergence that may reflect differences in the architecture of the resulting co-assemblies: RNA drives PSMα3 into structured α-helical fibrillar polymorphs, and LL-37 into disordered, amorphous co-aggregates.

Interestingly, although LL-37 does not adopt canonical amyloid architectures, it displays multiple functional connections to amyloid systems. LL-37 has been reported to bind the Alzheimer’s-associated amyloid-β (Aβ) peptide and inhibit its fibrillization, likely through stabilization of prefibrillar intermediates (De Lorenzi et al., 2017). This observation suggests that LL-37 can influence amyloid assembly pathways rather than simply interacting with mature fibrils, with potential implications for aging, immune regulation, and infection. In addition, LL-37 has been shown to interact with human α-synuclein as well as with bacterial curli amyloids, where it modulates biofilm properties (Kai-Larsen et al., 2010; Santos et al., 2023, 2021). Together, these findings position LL-37 as a cross-reactive regulator of amyloid assemblies across host and microbial contexts. The mechanistic basis and physiological relevance of these interactions remain to be further elucidated.

### Parallels with phase-transition biology

The RNA-dependent phase transitions observed for PSMα3 parallel broader principles established in phase-transition biology, where nucleic acids regulate the assembly, material properties, and function of biomolecular condensates. Similar RNA-driven liquid-to-solid transitions have been described for human RNA-binding proteins such as FUS and TDP-43, in which LLPS enables dynamic assembly states that can mature into more solid-like structures associated with pathology (Ambadipudi et al., 2017; Babinchak et al., 2019; Boyko and Surewicz, 2022; Elbaum-Garfinkle, 2019; Hyman et al., 2014; Kanaan et al., 2020; Mukherjee et al., 2024; Murakami et al., 2015; Patel et al., 2015; Ray et al., 2020; Röntgen et al., 2025; Wang et al., 2021; Y.-L. Wang et al., 2023; Wegmann et al., 2018). Antimicrobial peptides have likewise been shown to undergo nucleic-acid-dependent co-assembly and phase separation (Sneideris et al., 2023), indicating that RNA-mediated control of peptide assembly extends across structurally diverse systems.

Within this framework, the nucleolar accumulation of PSMα3 (Fig. 5) is notable because the nucleolus itself is an RNA-rich phase-separated compartment. The constrained but measurable fluorescence recovery of PSMα3 within nucleoli (Fig. S8) parallels the reduced dynamics observed during aging of RNA–PSMα3 condensates in vitro, although we do not infer a direct mechanistic equivalence between the cellular and in vitro states. Given the rapid membrane disruption induced by PSMα3 at cytotoxic concentrations (Figs. 5–6, and Movies S1–S2), we do not interpret nucleolar localization as evidence for a distinct intracellular toxic mechanism. Rather, it reflects the intrinsic nucleic-acid binding capacity of PSMα3 following cellular entry. Whether such interactions become biologically relevant at sub-cytotoxic concentrations, where membrane disruption does not dominate, remains an important question for future investigation.

### PSMα3 amyloid-like assembly at the intersection of bacterial virulence and human amyloid biology

The functional relevance of PSMα3 amyloid-like assembly extends beyond bacterial virulence to interactions with human amyloid systems, similarly to LL-37. Its effects are assembly state-dependent: monomeric α-helical PSMα3 inhibits fibrillation of insulin and Aβ40, whereas oligomeric and cross-α fibrillar forms promote Aβ40 aggregation (Kalitnik et al., 2025, 2024; Peng et al., 2023), indicating that supramolecular architecture governs its activity. PSMα3 also binds α-synuclein oligomers with low nanomolar affinity, targeting an oligomer-specific N-terminal motif and inhibiting oligomer-associated toxicity, with no interaction observed for monomeric α-synuclein (Santos et al., 2024, 2021). This selectivity highlights the emergence of functional surfaces unique to assembled states. Moreover, our data support a broader principle in which supramolecular architecture serves as a dynamic determinant of biological function. Structured fibrillar assemblies are required for LL-37 activity and membrane interactions (Sancho-Vaello et al., 2017), nanoscale organization controls β-defensin immunostimulatory potency (Lee et al., 2019; Schmidt et al., 2015; Tewary et al., 2013), and α-helical assembly states correlate with cytotoxicity across fibril-forming antimicrobial peptides (Bücker et al., 2022; Landau et al., 2026; Ragonis-Bachar et al., 2022; Salinas et al., 2021; Strati et al., 2025).

Together, these findings argue against PSMα3 assembly as incidental and instead support a model of functionally relevant, assembly-dependent activity. Notably, the RNA-dependent modulation of PSMα3 described here raises the possibility that extracellular nucleic acids in inflammatory environments could regulate its interactions with human amyloids, linking infection to neurodegenerative pathways.

### Biological and therapeutic implications

The divergent outcomes of RNA-mediated modulation, namely preservation of PSMα3 virulence versus attenuation of LL-37 cytotoxicity, reflect peptide-specific assembly architectures and suggest that RNA may tilt the host-pathogen balance toward bacterial advantage in infection-relevant environments rich in extracellular nucleic acids. Collectively, the findings position RNA as an environmental regulator of α-helical peptide assemblies, tuning bioactivity by modulating supramolecular organization and reversibility.

Nucleic acids are abundant in infection-relevant settings where PSMs operate, including biofilms enriched in extracellular DNA and RNA, abscesses, neutrophil extracellular traps, and damaged tissues that release host RNA. In such contexts, RNA could control the persistence and timing of membrane-active PSM species by shifting assemblies between condensate-like reservoirs and more inert aggregates, thereby modulating virulence without invoking an intracellular mechanism. This raises the possibility that extracellular RNA can influence the host–pathogen balance in inflammatory milieus by differentially regulating bacterial and host peptide assemblies.

While Poly (AU) served here as a defined, reductionist probe of peptide–RNA interactions, multiple endogenous RNA pools are plausible physiological partners. Within *S. aureus* biofilms, extracellular RNA is an abundant matrix component that contributes structurally to biofilm organization, raising the possibility that it directly modulates PSM assembly and thereby impacts biofilm stability and virulence-factor availability (Chiba et al., 2022; Mugunthan et al., 2023). Beyond biofilms, amyloid formation by PSMs has been detected in planktonic cultures, where δ-toxin–containing fibrils co-purify with extracellular membrane vesicles from virulent strains (X. Wang et al., 2023), linking amyloid formation to active secretion pathways. Conversely, the absence of detectable fibrils in certain in vivo infection models (X. Wang et al., 2023) may reflect the strong concentration dependence and environmental sensitivity of PSM assembly. Host-derived extracellular RNA released from stressed or dying cells can additionally function as a danger-associated molecular pattern (Preissner et al., 2020; Preissner and Fischer, 2023) and may further influence the balance between PSMα3-driven cytotoxicity and LL-37-mediated host protection. Defining which endogenous RNA species, concentrations, and structural features govern these assembly-dependent effects in vivo therefore represents an important next step.

More broadly, this work shifts the conceptual focus from amyloid formation as a static endpoint to environmentally regulated phase and assembly transitions. The findings further demonstrate that RNA-mediated regulation extends beyond canonical amyloids to structurally distinct α-helical peptide assemblies. For PSMα3, RNA-sensitive assembly dynamics may enable *S. aureus* to tune virulence in response to local environmental cues, supporting immune evasion, biofilm-associated persistence, and survival across diverse host niches. For LL-37, RNA-mediated attenuation of cytotoxicity without loss of antimicrobial activity suggests a host-protective mechanism that balances pathogen control with limiting collateral tissue damage. Together, these insights motivate therapeutic strategies aimed at reshaping assembly pathways and material states, rather than simply blocking aggregation.

## Materials and Methods

| Key Resources Table |  |  |  |  |
| --- | --- | --- | --- | --- |
| Reagent type (species) or resource | Designation | Source or reference | Identifiers | Additional information |
| Peptide | PSMα3 (Unlabeled or with FITC-labeled C-terminus) | Custom synthesis; GL Biochem (Shanghai) Ltd. | P0C805 · PSMA3_STAA8 | |
| Peptide | LL-37 (Unlabeled or with FITC-labeled C-terminus) | Custom synthesis; GL Biochem (Shanghai) Ltd. | P49913 · CAMP_HUMAN | LL-37 is generated by proteolytic cleavage of the cathelicidin antimicrobial peptide precursor (CAMP). |
| Sequence-based reagent | PolyA-IRDye 800CW and Poly (U) RNA | IDT |  |  |
| Sequence-based reagent | Poly(AU) RNA | Sigma Aldrich |  |  |
| Chemical compound | Thioflavin T (ThT) | Sigma Aldrich |  |  |
| Chemical compound | epigallocatechin gallate (EGCG) | Thermo Scientific Chemicals | Cat no. 449010100 |  |
| Cell line (Homo sapiens) | HeLa | ATCC® CCL-2™ |  | Human cervical epithelial cell line |
| Strain, strain background (*Escherichia coli*) | RFM795 | CGSC, Yale University (obtained as a kind gift from Prof. Sima Yaron, Technion) | CGSC# 14179 | Contains *lptD4213* deletion; increased outer membrane permeability |
| Commercial assay or kit | Lactate dehydrogenase (LDH) release assay | Roche Applied Science (Sigma-Aldrich) | Cat. no. 04744934001 | Cytotoxicity assays |
| Commercial assay or kit | PrestoBlue™ Cell Viability Reagent | Invitrogen (Thermo Scientific Chemicals) | Cat no. A13261 | Bacterial viability assay |
| Software | GraphPad Prism (v11) | GraphPad Software | RRID:SCR_002798 | statistical analysis |
| Software | Fiji/ImageJ | https://imagej.net/software/fiji/ | RRID:SCR_002285 | Image processing |
| Software | ZEISS ZEN | Carl Zeiss | RRID:SCR_013672 | TIRF and FRAP image acquisition and processing. |
| Software | Imaris (Oxford Instruments) | Oxford Instruments | RRID:SCR_007370 | Live-cell confocal movie analysis. |
| Software | TopSpin 4.1.4 | Bruker BioSpin | RRID:SCR_014227 | NMR data acquisition and processing |
| Software | POKY | Lee et al., 2021 (ref. 121) | https://poky.clas.ucdenver.edu | NMR spectral assignment and analysis |
| Software | Scilab | Scilab Enterprises | https://www.scilab.org/<br>RRID:SCR_014258 | NMR numerical computation, data analysis, and custom scripting |
| Major equipment | Transmission electron microscope | Talos F200C ThermoFisher Scientific (FEI) | Ilse Katz Institute for Nanoscale Science & Technology, Ben Gurion University of the Negev | TEM imaging of fibrils and aggregates |
| Major equipment | Spinning-disk confocal | Ti2-E + CSU-W1 Nikon / Yokogawa | LS&E Infrastructure Center, Technion–Israel Institute of Technology | Live-cell imaging of PSMα3 in HeLa cells |
| Major equipment | Laser-scanning confocal | LSM 710 Carl Zeiss | LS&E Infrastructure Center, Technion–Israel Institute of Technology | FRAP (in vitro condensates & nucleolar) |
| Major equipment | Widefield fluorescence microscope | DMI8 Leica | LS&E Infrastructure Center, Technion–Israel Institute of Technology | Fluorescence imaging of peptide–RNA mixtures in vitro |
| Major equipment | Super-resolution / TIRF microscope | Elyra 7 Carl Zeiss | LS&E Infrastructure Center, Technion–Israel Institute of Technology | TIRF (AmyTracker), |
| Major equipment | NMR spectrometer | 700 MHz AVANCE NEO + Prodigy cryoprobe Bruker BioSpin | Max Planck Institute for Multidisciplinary Sciences, Göttingen | 1D <sup>1</sup>H, 2D TOCSY & NOESY of PSMα3–EGCG |
| Major equipment | Multimode plate reader | FLUOstar Omega BMG Labtech | Technion–Israel Institute of Technology | ThT fibrillation kinetics, turbidity (OD<sub>400</sub>), LDH absorbance |
| Major equipment | Multimode plate reader | CLARIOstar<br>BMG Labtech | Technion–<br>Israel Institute<br>of Technology | PrestoBlue<br>bacterial<br>viability assay |
| Major equipment | CD<br>spectropolarimeter | Chirascan (with<br>solid sample<br>holder<br>CS/SSH)<br>Applied<br>Photophysics | SPC facility,<br>EMBL<br>Hamburg | Solid-state CD<br>of PSMα3<br>secondary<br>structure |
| Major equipment | Gel imaging<br>system | Odyssey FC<br>LI-COR | Technion–<br>Israel Institute<br>of Technology | EMSA<br>visualization<br>(IRDye 800CW<br>channel) |

### Peptide and Poly (AU) RNA preparation

PSMα3 (sequence: MEFVAKLFKFFKDLLGKFLGNN) and LL-37 (sequence: LLGDFFRKSKEKIGKEFKRIVQRIKDFLRNLVPRTES), either unlabeled, or with a C-terminal FITC-labeled (PSMα3-FITC and LL-37-FITC) were purchased from GL Biochem (Shanghai) Ltd as custom synthesis with >98% purity. Poly (AU) RNA was purchased from Sigma Aldrich. A stock solution of the peptides was prepared at a concentration of 1 mM in a mixture of 20% dimethyl sulfoxide (DMSO) and 80% ultra-pure water. For light microscopy experiments, we used a mixture of 20% FITC-labeled and 80% unlabeled PSMα3 or LL-37. The Poly (AU) RNA stock solution was prepared at a concentration of 2000 ng/µl in UPW, the stock solution was stored at −80 °C until further use. Labeled Poly (AU) RNA was prepared by introducing 0.02 mg/ml of PI dye to the Poly (AU) RNA stock solution. For the experiments, samples containing 100 µM PSMα3 or LL-37 with Poly (AU) RNA at 10, 50, 100, 200, or 400 ng/µl were prepared in 50 mM HEPES buffer with 150 mM sodium chloride (NaCl), adjusted to pH 7.4, or in 20 mM Tris, 20 mM Bis-Tris, and 20 mM sodium acetate adjusted to pH 4.

### Cytotoxicity against HeLa cells tested using the lactate dehydrogenase (LDH) release assay

Human cervical carcinoma HeLa cells (ATCC® CCL-2™) were routinely cultured in Dulbecco’s Modified Eagle’s Medium - high glucose (DMEM) (Sigma, Israel) with L-glutamine and supplemented with penicillin (100 U/ml), streptomycin (0.1 mg/ml), and with 10% fetal calf serum (Sigma, Israel). The cells were grown at 37°C and 5% CO_2_. One day before the experiment, cells were resuspended in growing medium (DMEM supplemented with 10% fetal calf serum) 1×10^5^ cells/ml. 50 μL of cell suspension were pipetted into a 96-well plate and grown over night. Thirty minutes before the experiment cells were washed and resuspended in 50 μL of DMEM medium supplemented with L-glutamine (100 U/mL), and with penicillin (100 U/mL), streptomycin (0.1 mg/mL), and with 0.5% heat-inactivated fetal calf serum (assay medium). For the cytotoxicity assay, 100 µM PSMα3 was incubated with or without EGCG at molar ratios of 1:1 or 1:5, or with Poly (AU) RNA at concentrations of 50 ng/µL or 400 ng/µL, for the designated incubation times. Similarly, 100 µM LL-37 was prepared under the same conditions, with or without EGCG at 1:1 or 1:5 molar ratios, or Poly (AU) RNA at 100 ng/µL or 400 ng/µL, following the specified incubation times.

Serial two-fold dilutions in assay medium were performed, and 50 μL of each dilution were pipetted into the three different 96-well with the cells. The plates were incubated for 30 min at 37°C and 5% CO2 and then cell lysis was quantified using the LDH release colorimetric assay according to the manufacturer’s instructions, including all recommended controls (LDH; Cytotoxicity Detection Kit Plus, Roche Applied Science, Germany). Cell-free assay medium was measured as background. Cells subjected to the same experimental conditions apart from peptide addition were used as a control to account for spontaneous LDH release. Cells subjected to the same experimental conditions apart from peptide addition and treated with manufacturer-supplied lysis buffer were used as a control to account for maximum LDH release. Absorbance at 490 and 690 nm was measured in a plate reader (FLUOstar Omega, BMG Labtech, Germany). Absorbance at 690 nm was subtracted from 490 nm readings to correct for background. The mean absorbance of triplicate samples and controls was calculated, followed by background subtraction. Data was obtained from at least three independent biological replicates, with the arithmetic mean used for averaging. Error bars represent the standard deviation (SD). Statistical analysis, including one-way ANOVA, was performed using GraphPad Prism 11.

### Assessment of Bacterial Viability Using PrestoBlue™ HS Cell Viability Reagent

*Escherichia coli (E. coli)* strain RFM795 (*lptD4213*), obtained from the *E. coli* Genetic Stock Center (CGSC#14179, Yale University) as a kind gift from Prof. Sima Yaron (Technion – Israel Institute of Technology), was used for all antibacterial activity assays. This strain carries an in-frame deletion in *lptD* (residues D330–D352), resulting in increased outer membrane permeability that enhances peptide access to the bacterial membrane. Bacterial cultures were grown overnight in Luria Broth (LB) medium at 37 °C with shaking at 220 rpm. PSMα3 and LL-37 stock solutions were prepared by dissolving the peptides at a concentration of 1 mM in a solvent mixture of 20% DMSO and 80% ultra-pure water. From these stock solutions, the following sample preparations were made: For PSMα3: **(1)**100 µM PSMα3 **(2)**100 µM PSMα3 with 50 ng/µL Poly (AU) RNA **(3)** 100 µM PSMα3 with 400 ng/µL Poly (AU) RNA **(4)** 100 µM PSMα3 with 500 µM EGCG. For LL-37: **(1)**100 µM LL-37 **(2)**100 µM LL-37 with 100 ng/µL Poly (AU) RNA **(3)** 100 µM LL-37 with 400 ng/µL Poly (AU) RNA **(4)** 100 µM LL-37 with 500 µM EGCG.

All solutions were prepared in 50 mM HEPES buffer pH 7.4 containing 150 mM NaCl. Where applicable, samples were incubated for the specified durations of 2 hours or 24 hours.

On the day of the experiment, dilutions of the PSMα3 solutions to 20 µM and 10 µM were prepared in LB medium and dispensed into a sterile 96-well black flat-bottom plate (Greiner bio-one). *E. coli* cultures were diluted to an optical density (OD) of 0.2, and bacterial suspensions were added to the wells. The plate was incubated at 37 °C with shaking at 220 rpm for 1 hour to allow the reaction to proceed.

Following incubation, 10X PrestoBlue™ HS Cell Viability Reagent (Invitrogen) was added to the wells. Wells containing only LB medium served as negative controls, while wells containing only *E. coli* served as positive controls. Plates were sealed with a thermal seal film (EXCEL Scientific) and incubated in a plate reader (CLARIOstar). Bacterial viability was assessed using PrestoBlue fluorescence (excitation: 535–560 nm; emission: 590–615 nm) over time. Each condition was tested in triplicate across three independent experiments. Fluorescence values were averaged, and blank readings were subtracted. Antimicrobial activity was determined at the 2-hour time point, when fluorescence was highest in the positive control. Data were obtained from at least three independent biological replicates, with the arithmetic mean used for averaging. Error bars represent the standard deviation (SD). Statistical analysis, including one-way ANOVA, was performed using GraphPad Prism 11.

### Transmission electron microscopy (TEM)

For TEM analysis, 4–5 µL of 100 µM PSMα3, with or without EGCG or Poly (AU) RNA at concentrations of 10 ng/µL, 50 ng/µL, and 400 ng/µL, as well as 100 µM LL-37, with or without Poly (AU) RNA at 100 ng/µL and 400 ng/µL, were directly applied onto glow-discharged 400-mesh copper grids (easiGlow; Pelco, Clovis, CA, USA) with a grid hole size of 42 µm, stabilized with Formvar/carbon (Ted Pella, Inc.). The grids were glow-discharged using a 15-mA current with a negative charge for 25 seconds. Samples were allowed to adhere to the grids for 45 seconds before being stained with a 1% uranyl acetate solution (Electron Microscopy Science, 22400-1) for 45 seconds. The samples were then examined using a ThermoFisher Scientific (FEI) Talos F200C transmission electron microscope, operating at 200 kV and equipped with a Ceta 16M CMOS camera, at the Ilse Katz Institute for Nanoscale Science and Technology, Ben Gurion University of the Negev, Israel.

### Turbidity measurements

Turbidity measurements were performed using a FLUOstar Omega plate reader (BMG Labtech) set to a wavelength of 400 nm. For each sample, the protein concentration was maintained at 100 µM, while RNA concentrations were varied across the following levels: 10 ng/µl, 20 ng/µl, 50 ng/µl, 100 ng /µl, 200 ng/µl, and 400 ng/µl. The turbidity of the mixtures was monitored over several hours following resuspension. The maximum absorbance recorded within the first 30 minutes was used as an indicator of the dense phase volume. The experiments were conducted three times with similar observations.

### Electrophoretic mobility shift assay (EMSA)

RNA molecules were synthesized by IDT, with oligoA modified by the IRDye 800CW fluorescent dye. Double-stranded RNA was prepared by annealing 10 µM oligoA-IRDye 800CW with 10 µM oligoU in an annealing buffer containing 10 mM Tris-HCl (pH 7.5), 50 mM NaCl, and 1 mM EDTA. The mixture was heated to 95°C for 5 minutes, followed by gradual cooling to 25°C at a rate of 2°C per minute. The final RNA concentration used in the binding reactions was 40 nM (∼400ng/µL).

RNA-protein complexes were incubated at 37°C for 30 minutes before being resolved on a 2.5% agarose gel. Gel electrophoresis was performed in a 0.5× TBE buffer for 7 minutes. The complexes were visualized using a Li-Cor Odyssey FC imaging system in the 800 nm channel with a 30-second exposure. For the EMSA, binding reactions were carried out in a buffer composed of 20 mM HEPES (pH 7.9), 50 mM KCl, 1 mM Dithiothreitol (DTT), 0.1 mM Ethylenediaminetetraacetic acid (EDTA), 5% glycerol, and 0.05% nonyl phenoxypolyethoxylethanol (NP-40). PSMα3 was added to the reactions at final concentrations of 0, 40, 80, 160, and 320 µM. The experiments were conducted three times with similar observations.

### Fluorescent microscopy imaging of peptides with RNA in vitro

Prior to imaging, 100 µM PSMα3 or LL-37 (containing 20% FITC-labeled and 80% unlabeled peptide) were combined with PI-labeled Poly (AU) RNA at different concentrations in 50 mM HEPES buffer pH 7.4 containing 150 mM NaCl. Sample were also tested after 65°C heat shock for 15 minutes. In addition, the PSMα3 and RNA mixture was also tested in a different buffer containing 20 μM Tris, 20 μM Bis-Tris, and 20 μM sodium acetate at pH 4 and were subjected to heat shock at 65°C for 15 minutes.

Following preparation, with or without heat shock, a 10 µL aliquot of each sample was transferred to a µ-Slide 8 Well ibidi chamber or to a clean 24 × 60 mm No. 1.5 glass slide for imaging. Fluorescence microscopy was performed using a Leica DMI8 inverted fluorescent microscope equipped with a 63× immersion oil objective (Numerical Aperture, NA = 1.4) at the Life Sciences and Engineering (LS&E) Infrastructure Center, Technion-Israel Institute of Technology, Haifa, Israel. Data were processed and analyzed using Fiji-ImageJ software. The experiment was conducted three times with similar observations.

### Fluorescence recovery after photobleaching (FRAP) measurements

FRAP experiments were conducted using a Zeiss LSM 710 laser scanning confocal microscope equipped with a 63× Plan-Apochromat oil immersion objective, NA1.4, at the Life Sciences and Engineering (LS&E) Infrastructure Center, Technion-Israel Institute of Technology, Haifa, Israel. The experiments were performed on condensates from 100 µM PSMα3 or LL-37 (containing 20% FITC-labeled and 80% unlabeled peptide) mixed with PI-labeled Poly (AU) RNA at various concentrations as indicated in the figures. Photobleaching was performed using 405 nm and 488 nm laser lines. The experiments were conducted three times with similar observations.

For FRAP analysis inside the nucleolus of HeLa cells, PSMα3 was added to the cells at a final concentration of 20 µM containing 20% FITC-labeled and 80% unlabeled peptide. The cells were incubated with the peptide for 20 minutes before conducting the experiment. Data was processed and analyzed using ZEISS ZEN software and Fiji-ImageJ software. The experiments were conducted three times with similar observations.

### Total internal reflection fluorescence (TIRF) microscopy

TIRF microscopy was performed using a ZEISS Elyra 7 Super-Resolution Microscope equipped with a 63× Apochromat alpha Plan-Apochromat Oil immersion DIC objective, NA1.46, at the Life Sciences and Engineering (LS&E) Infrastructure Center, Technion-Israel Institute of Technology, Haifa, Israel. Images were acquired using the pco.edge sCMOS cameras, and data were processed and analyzed with ZEISS ZEN software to accurately represent the observed phenomena. The condensates and aggregates analyzed were generated using 100 µM of 20% FITC-PSMα3 (green) and 80% unlabeled PSMα3 with either 50 ng/µL or 400 ng/µL Poly (AU) RNA. AmyTracker 630 (Ebba Biotech) was added at a 1:500 (peptide: AmyTracker) molar ratio to each sample. The experiments were conducted three times with similar observations.

### Confocal microscopy visualization of PSMα3 interaction with HeLa cells

HeLa cells were pre-cultured one day before the experiment by preparing a suspension containing 350,000 cells/ml and plating 150 µl of this suspension into each well of a µ-Slide 8 well glass-bottom chamber. The cells were then incubated overnight under standard growth conditions (37°C, 5% CO2) to allow for adherence and growth. On the day of the experiment, the cells were washed three times with phosphate-buffered saline (PBS) to remove any residual media. Hoechst 33342 dye (10 mg/mL stock) was diluted 1:2000 in fresh cell media and added to the cells. The cells were incubated with Hoechst for 10 minutes at 37°C and 5% CO2. After incubation, cells were washed three times with PBS to remove Hoechst residuals. A working solution of PI was prepared by diluting a 1 mg/ml PI stock solution to a final concentration of 0.02 mg/ml in fresh cell growth media. Immediately prior to imaging, PSMα3 was added to the cells at a final concentration of 20 µM containing 20% FITC-labeled and 80% unlabeled peptide. The cells were then imaged using a Ti2-E microscope by Nikon with a CSU-W1 spinning disk confocal unit by Yokogawa, equipped with a 100X CFI SR HP Plan Apochromat Lambda S silicone immersion objective, NA1.35, at the Life Sciences and Engineering (LS&E) Infrastructure Center, Technion-Israel Institute of Technology, Haifa, Israel. Confocal movies and images were captured by Photometrics BSI sCMOS cameras to observe the interaction and localization of PSMα3 within the cells, focusing particularly on its colocalization with nucleic acids in the nucleolar region. The acquired images and movies were subsequently analyzed using Imaris Image Analysis Software (Oxford Instruments). The experiments were conducted three times with similar observations.

### Super resolution light microscopy visualization of PSMα3 interaction with *E. coli*

*E. coli* inoculum was pre-cultured in Luria-Bertani medium (LB) medium at 220 rpm and 37°C for 24 hours prior to the experiment. On the day of the experiment, the optical density (OD) of the culture was measured, and the cells were diluted to an OD of 0.4. The cells were then incubated with 20 μM of 20% FITC-PSMα3 (green) and 80% unlabeled PSMα3, with and without EGCG at 1:5 molar ratio, alongside a bacteria-only control. Samples were incubated for 30 minutes at 220 rpm and 37°C. Following incubation, the samples were centrifuged three times for 3 minutes at 1.5g, with the supernatant discarded and the pellet resuspended in 1X PBS after each spin. The resuspended cells were then stained with 50 μg/mL wheat germ agglutinin (WGA) CF633 for 30 minutes. After staining, the cells were washed three times by centrifugation with PBS to remove excess WGA. During the final wash, the cells were resuspended in 0.1 mg/mL PI solution.

Prior to imaging, the samples were loaded into an ibidi µ-Slide VI 0.4. Imaging was performed using a ZEISS Elyra 7 Super-Resolution Microscope equipped with a 63× Apochromat alpha Plan-Apochromat Oil immersion DIC objective, NA 1.4, and pco.edge sCMOS cameras. Lattice structured illumination images were acquired and processed using the ZEN black software at the Life Sciences and Engineering (LS&E) Infrastructure Center, Technion-Israel Institute of Technology, Haifa, Israel. The experiments were conducted three times with similar observations.

### Thioflavin T fluorescence fibrillation kinetics assay

Thioflavin T (ThT) (Sigma Aldrich) is a widely used fluorescent dye for detecting and analyzing amyloid fibril formation kinetics. Fibrillation curves in the presence of ThT typically exhibit an initial lag phase followed by rapid fibril elongation. To accurately capture the fibrillation lag time, PSMα3 was pre-treated before the experiment. The peptide was dissolved in a 1:1 mixture of 1,1,1,3,3,3-Hexafluoroisopropanol (HFIP- Sigma Aldrich) and Trifluoroacetic acid (TFA- Sigma Aldrich) to a final concentration of 1 mg/mL, followed by bath sonication for 10 minutes at room temperature. The organic solvents were then evaporated using a mini-rotational vacuum concentrator (Christ, Germany) at 1,000 rpm for 2 hours at room temperature.

For the experiment, freshly prepared 100 µM PSMα3 peptides, with or without 500 µM EGCG at a 1:5 molar ratio, were prepared in 50 mM HEPES buffer pH 7.4 containing 150 mM NaCl. ThT was prepared by diluting a stock solution in ultrapure double-distilled water (UPddw) and filtered before use to reach a final concentration of 200 µM. Blank control solutions containing all components except for the peptide were prepared for each reaction.

The assay was conducted in black 96-well flat-bottom plates (Greiner Bio-One), which were sealed with a thermal seal film (EXCEL Scientific) to prevent evaporation. Samples were incubated in a plate reader (OMEGA) at 37°C, shaking at 500 rpm for 85 seconds before each reading cycle, for up to 1,000 cycles of 6 minutes each, totaling approximately 100 hours. Fluorescence was measured in triplicate using an excitation wavelength of 438 ± 20 nm and an emission wavelength of 490 ± 20 nm. All values were averaged, background fluorescence was subtracted using blank controls, and the results were plotted over time. Standard deviation is shown as error bars. The entire experiment was independently repeated at least three times on different days to ensure reproducibility.

### Solid-State Circular Dichroism Spectroscopy

Solid-state circular dichroism spectroscopy(ss-CD) was conducted with PSMα3 to assess its fibrillar secondary structure components in presence and absence of Poly (AU) RNA. For preparation of peptide and RNA samples, see Peptide and Poly (AU) preparation section. PSMα3 was prepared at a working concentration of 100 µM in 50 mM HEPES, pH 7.4, 150 mM NaCl at 0 ng/µL, 50 ng/µL or 400 ng/µL Poly (AU) RNA. The reaction mixes were incubated for 0 min, 2 h and 72 h at 37 °C in a non-shaking thermocycler. Following incubation, soluble reaction components were removed via dilution washing with 150 µL water and centrifugation at 12.500 rcf for 30 min. 20 µL of the sediment were resuspended in 180 µL water for a second washing centrifugation step at 12.500 rcf for 30 min. 180 µL of the supernatant were discarded and 18 µL of sediment applied to a *Chirascan Series fused silica disc* (AP/CSSD, Applied Photophysics) in three times 6 µL steps. After every sample application onto to the silica disc, all liquid was evaporated at 37 °C for 5–10 min until only an opaque film remained visible. For secondary structure determination of aggregated/fibrillar peptide components on the silica disc, the disc was inserted into a *Chirascan solid sample holder* (CS/SSH, Applied Photophysics) and the circular dichroism recorded in the far-UV (180–250 nm, step size 1 nm, time-per-point 1 s) using a CD spectropolarimeter (Applied Photophysics) at the Sample Preparation and Characterisation (SPC) facility at EMBL Hamburg located at the CSSB. To counteract sample anisotropy, CD spectra were recorded at least four different disc rotations and the results averaged. Prior to data analysis, silica disc and buffer/water backgrounds were subtracted from recorded sample spectra. Poly (AU) RNA-only background spectra showed complex and high amplitude signals that did not appear for samples in combination with peptides and were therefore not subtracted from sample spectra.

### NMR experiments

To prepare the reference samples for NMR analysis, 0.521 mg of PSMα3 was dissolved in 180 µL of 20 mM MES (2-(N-morpholino)ethanesulfonic acid) buffer at pH 6.0, supplemented with 5% (v/v) deuterium oxide (D₂O). From this solution, 170 µL was transferred into a 3.0 mm NMR tube for spectral acquisition. In parallel, a 4 mM stock solution of EGCG was prepared in the same 20 mM MES buffer (pH 6.0). From this, a working solution containing 0.5 mM EGCG in 20 mM MES with 5% (v/v) D₂O was prepared to match the buffer conditions of the peptide sample. A volume of 170 µL was then transferred into a separate 3.0 mm NMR tube for use as the EGCG reference.

All NMR experiments were recorded on a Bruker 700 MHz spectrometer equipped with a triple-resonance cryoprobe (Prodigy) and an AVANCE NEO console. One-dimensional (1D) ¹H-NMR spectra were acquired using the Bruker library pulse sequence zggpw5 for water suppression. The acquisition parameters were: acquisition time (AQ) = 1.90 s, spectral width (SW) = 12.31 ppm, recovery delay (D1) = 1.5 s, and 64 scans. All experiments were conducted at 35 °C, under which optimal spectral quality was obtained.

Two-dimensional (2D) ¹H–¹H TOCSY (Total Correlation Spectroscopy) spectra were collected using the Bruker pulse sequence dipsi2gpph19. Acquisition parameters were: *t*₂ = 2048 complex points (AQ₂ = 0.118 s), *t*₁ = 512 complex points (AQ₁ = 0.029 s), SW₁ = SW₂ = 12.31 ppm, with 64 scans and a mixing time (tmix) of 80 ms. Recovery delay (D1) was set to 1.5 s. 2D ¹H–¹H NOESY (Nuclear Overhauser Effect Spectroscopy) spectra were acquired using the noesyfpgpph19 pulse sequence, with acquisition parameters identical to those used for TOCSY. The mixing time was set to 300 ms. Total experimental time for the combined 2D TOCSY and NOESY experiments was approximately 30 hours. NMR data processing was carried out using TopSpin 4.1.4 (Bruker), and spectral assignments for the 2D TOCSY and NOESY spectra were performed using POKY software (Lee et al., 2021). Signal integration was carried out using in-house scripts written in Scilab 2025.0.0 (Campbell et al., 2010).

## Data availability

Raw data is also deposited in: https://doi.org/10.5281/zenodo.17598867

## Acknowledgements

M.L. acknowledges research support from the Israel Science Foundation, Grant No. 2111/20 and the Cure Alzheimer’s Fund. M.L. and M.Z. acknowledge the Forschungskooperation Niedersachsen – Israel, Volkswagenstiftung, No: 76251-4659/2022 (ZN 4042). Funded/Co-funded by the European Union (ERC, FuncAmyloid, 101087140). Views and opinions expressed are however those of the author(s) only and do not necessarily reflect those of the European Union or the European Research Council. Neither the European Union nor the granting authority can be held responsible for them. A.K.B thanks the Novo Nordisk Foundation (Grant number NNFSA17002839) and the European Union (ERC CoG 101088163 EMMA) for funding. We thank Ayala Shiber for advice and guidance, as well as Yael Mandel-Gutfreund and Hiba Hassanain for their assistance with RNA preparations. We acknowledge guidance and support from Nitsan Dahan from the Microscopy core facility center at the Lorry I. Lokey Interdisciplinary Center for Life Sciences and Engineering, and the technical support by the SPC facility at EMBL Hamburg. We acknowledge the use of OpenAI’s GPT-4, Claude and other AI-based models to improve the writing quality of parts of this manuscript.

---

**Figure S1.**
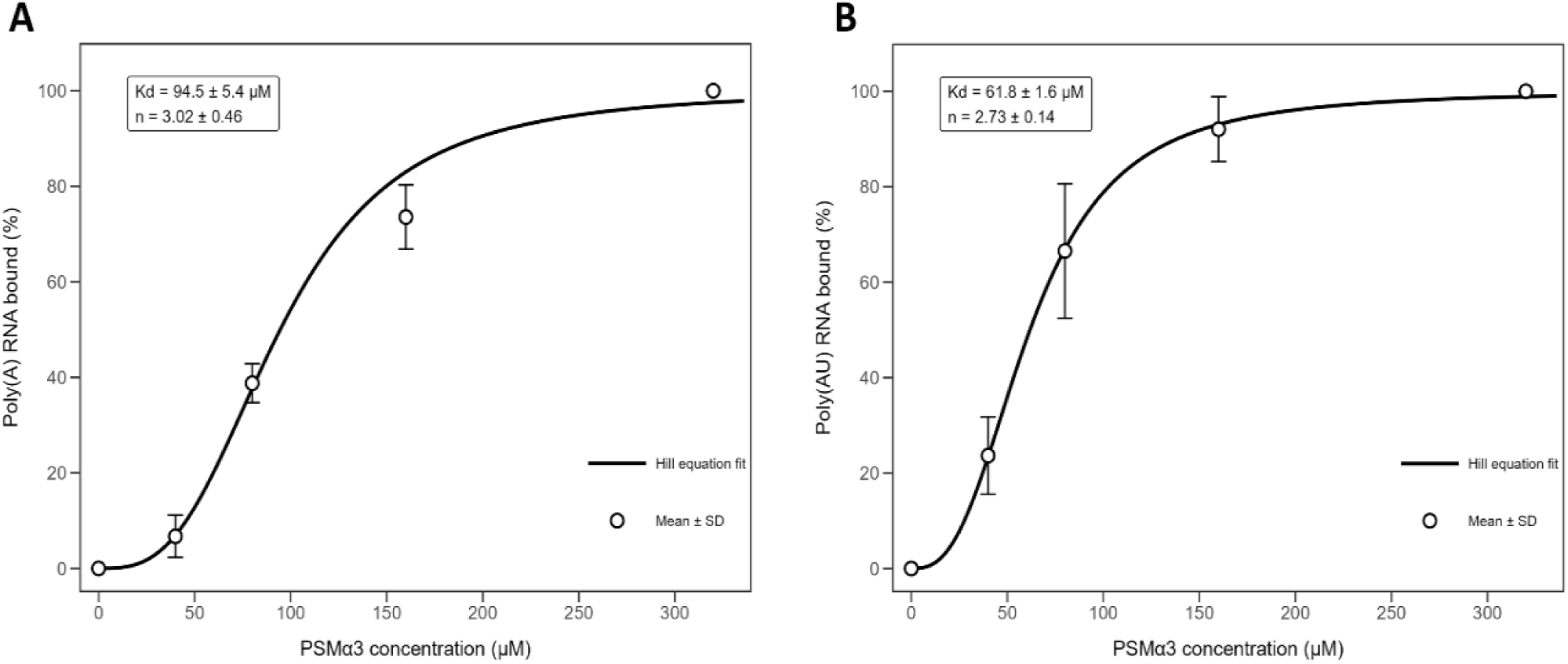
Concentration-dependent binding of PSMα3 to Poly (AU) and Poly (A) RNA. Percent RNA bound was quantified as a function of PSMα3 concentration for Poly (AU) **(A)** and Poly (A) **(B)** calculated from the EMSA gels shown in Figure 1. Data points represent mean ± SD. Solid lines indicate fits to a Hill binding model with fixed asymptotes (0–100%). The apparent dissociation constant (Kd) corresponds to the protein concentration yielding 50% RNA binding, fitted Kd and Hill coefficient (n) values ± SE are shown.

**Figure S2.**
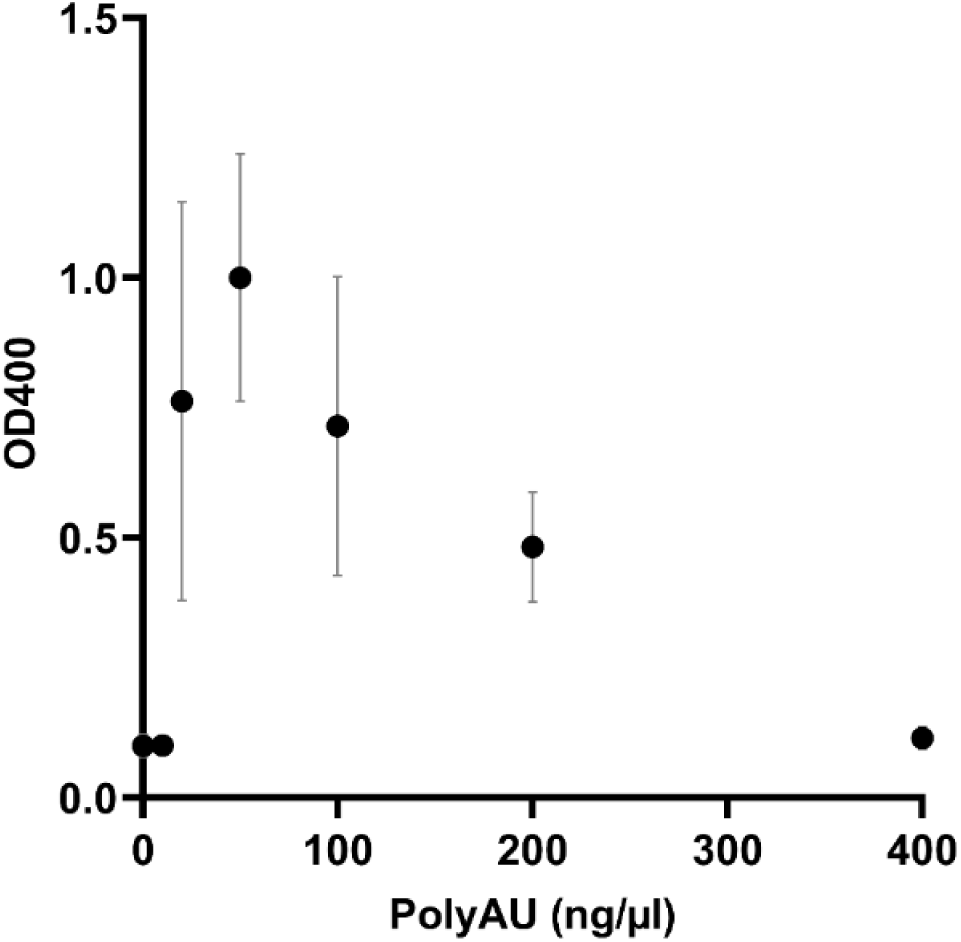
Turbidity measurements of PSMα3 with increasing Poly (AU) RNA concentrations. PSMα3 concentration was maintained at 100 µM, and Poly (AU) RNA concentrations varied across 0, 10, 20, 50, 100, 200, and 400 ng/µL. Data points represent the average optical density at 400 nm (OD400) recorded within the first 30 minutes after sample preparation, with each condition tested in triplicate. Error bars indicate the standard deviation.

**Figure S3.**
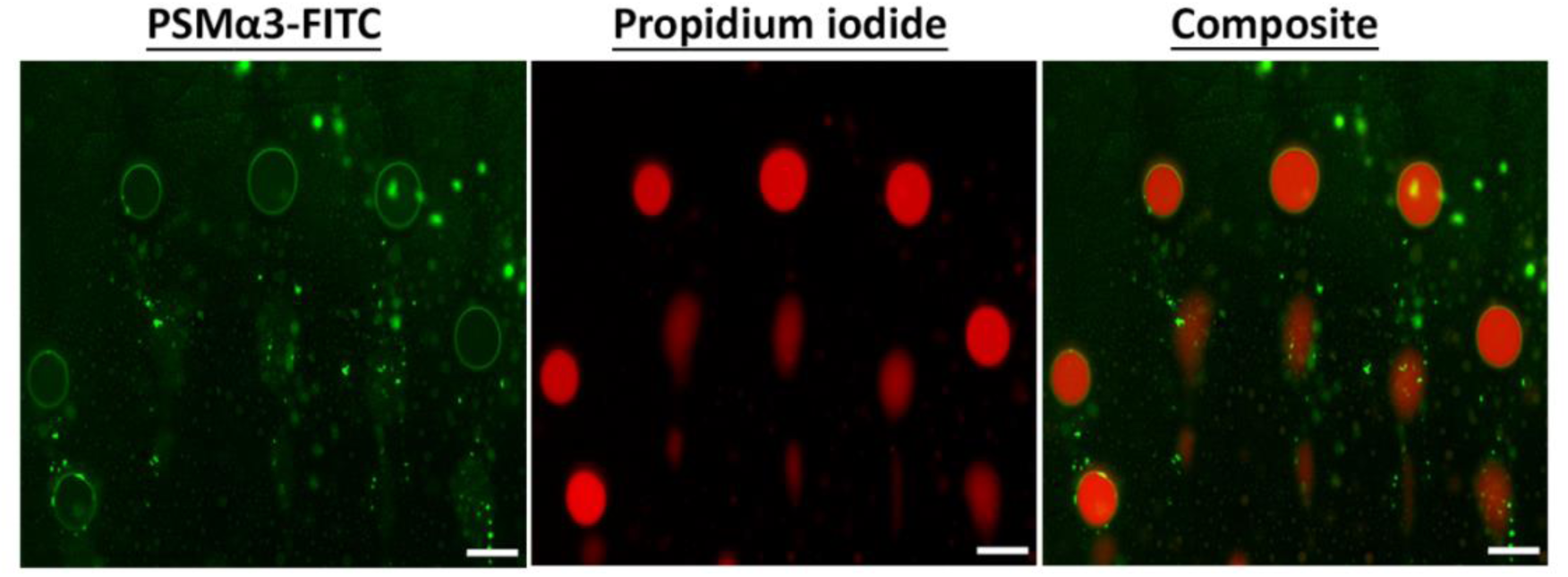
Encapsulation of Poly (AU) RNA within FITC-PSMα3 droplets. 20% FITC-PSMα3 (green) and 80% unlabeled 100 µM PSMα3 were mixed with 50 ng/µL Poly (AU) RNA labeled with propidium iodide (PI). The left panel shows the FITC green channel, the middle panel displays the PI red channel, and the right panel is a composite image showing the overlap of FITC-PSMα3 and Poly (AU) RNA-PI signals. Scale bars represent 20 µm.

**Figure S4.**
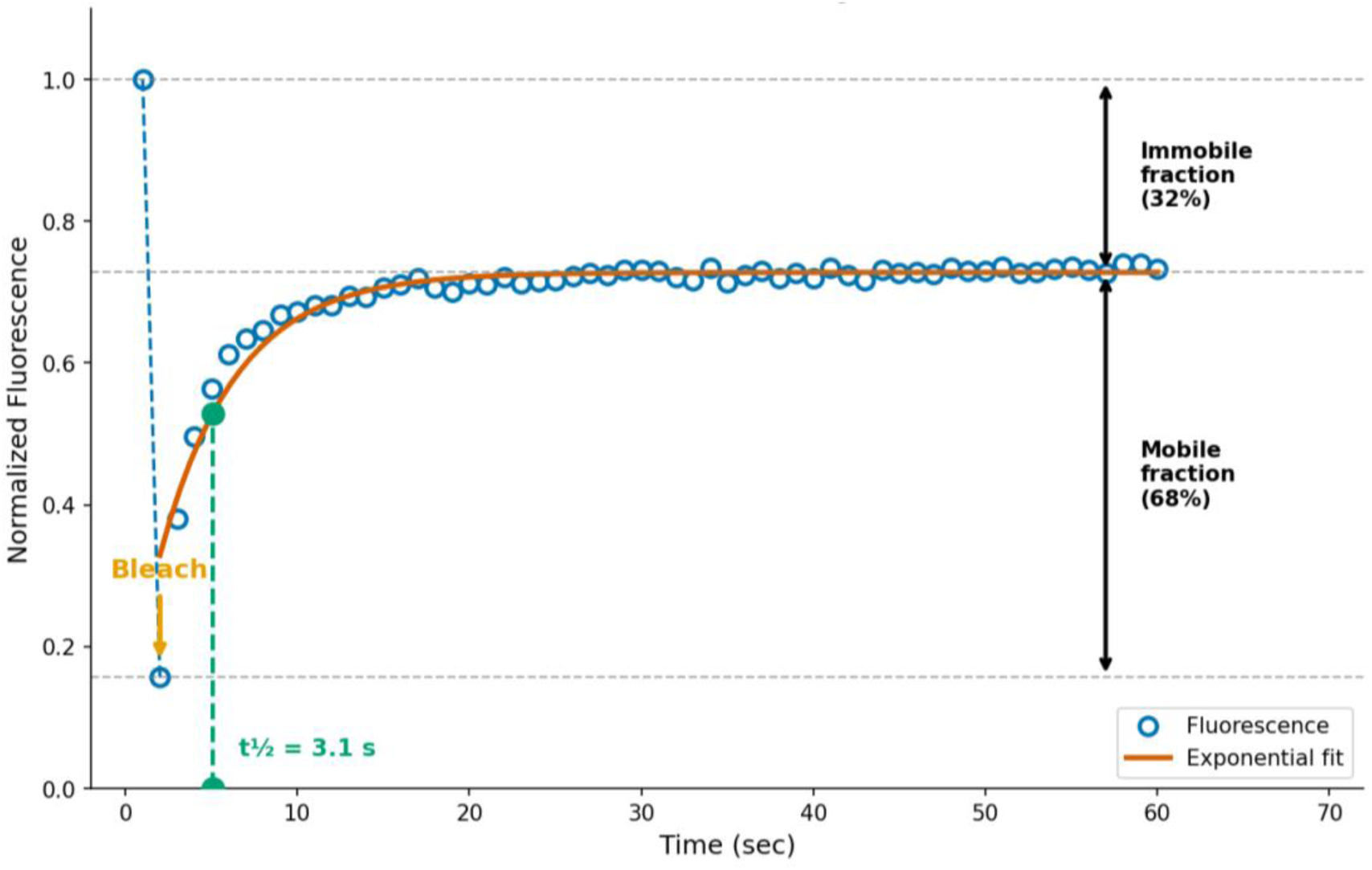
FRAP analysis of PSMα3 – Poly (AU) RNA condensate dynamics. Normalized fluorescence recovery after photobleaching (FRAP) measured for condensates formed by 20% FITC-PSMα3 (green) and 80% unlabeled 100 µM PSMα3 in the presence of 50 ng/μL Poly (AU) RNA, corresponding to the experiment shown in Figure 2. Blue circles represent measured fluorescence intensity normalized to pre-bleach levels, and the orange curve indicates a single-exponential fit to the recovery phase. The bleach event is marked by the orange arrow. Recovery reaches a plateau corresponding to a mobile fraction of 68% and an immobile fraction of 32%. The green marker and dashed line denote the half-time of recovery (t½ = 3.1 s). Horizontal dashed lines indicate pre-bleach intensity (top), plateau recovery level (middle), and immediate post-bleach intensity (bottom).

**Figure S5.**
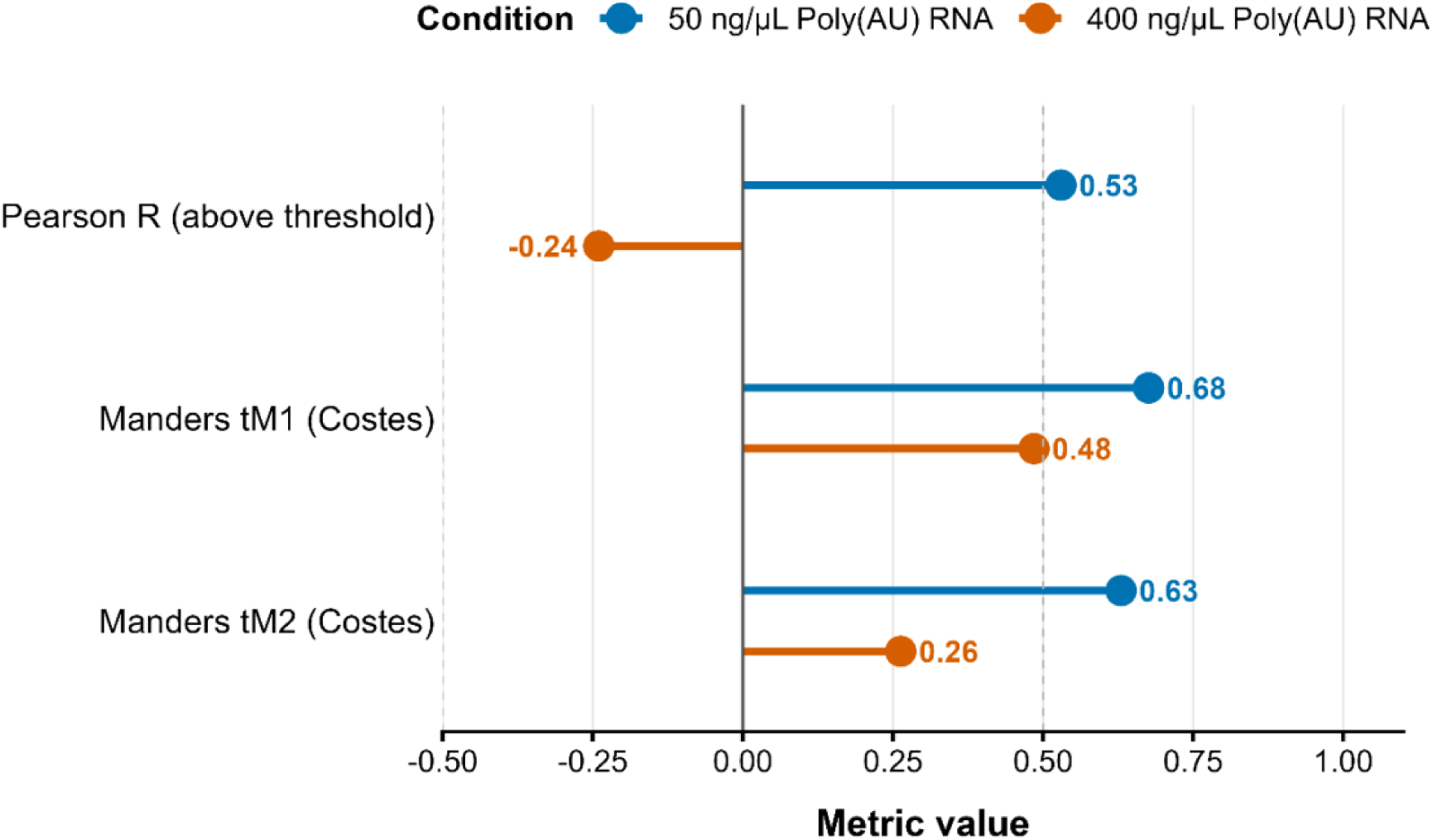
Colocalization analysis of PSMα3 and Poly (AU) RNA at different RNA concentrations. Quantitative comparison of colocalization metrics for 100 µM PSMα3 (20% FITC-PSMα3 and 80% unlabeled) and Poly (AU) RNA at 50 ng/μL (blue) and 400 ng/μL (orange), corresponding to the experiment shown in Figure 2. Pearson’s correlation coefficient was calculated using above-threshold pixels only, with thresholds determined automatically by the Costes method. Additional metrics include Spearman’s rank correlation coefficient (ρ), Li’s intensity correlation quotient (ICQ), and Manders’ colocalization coefficients (tM1 and tM2) calculated using Costes automatic thresholding. Numerical values for each metric are indicated on the plot. The dashed vertical line at 0.5 marks a commonly used reference value for strong colocalization.

**Figure S6.**
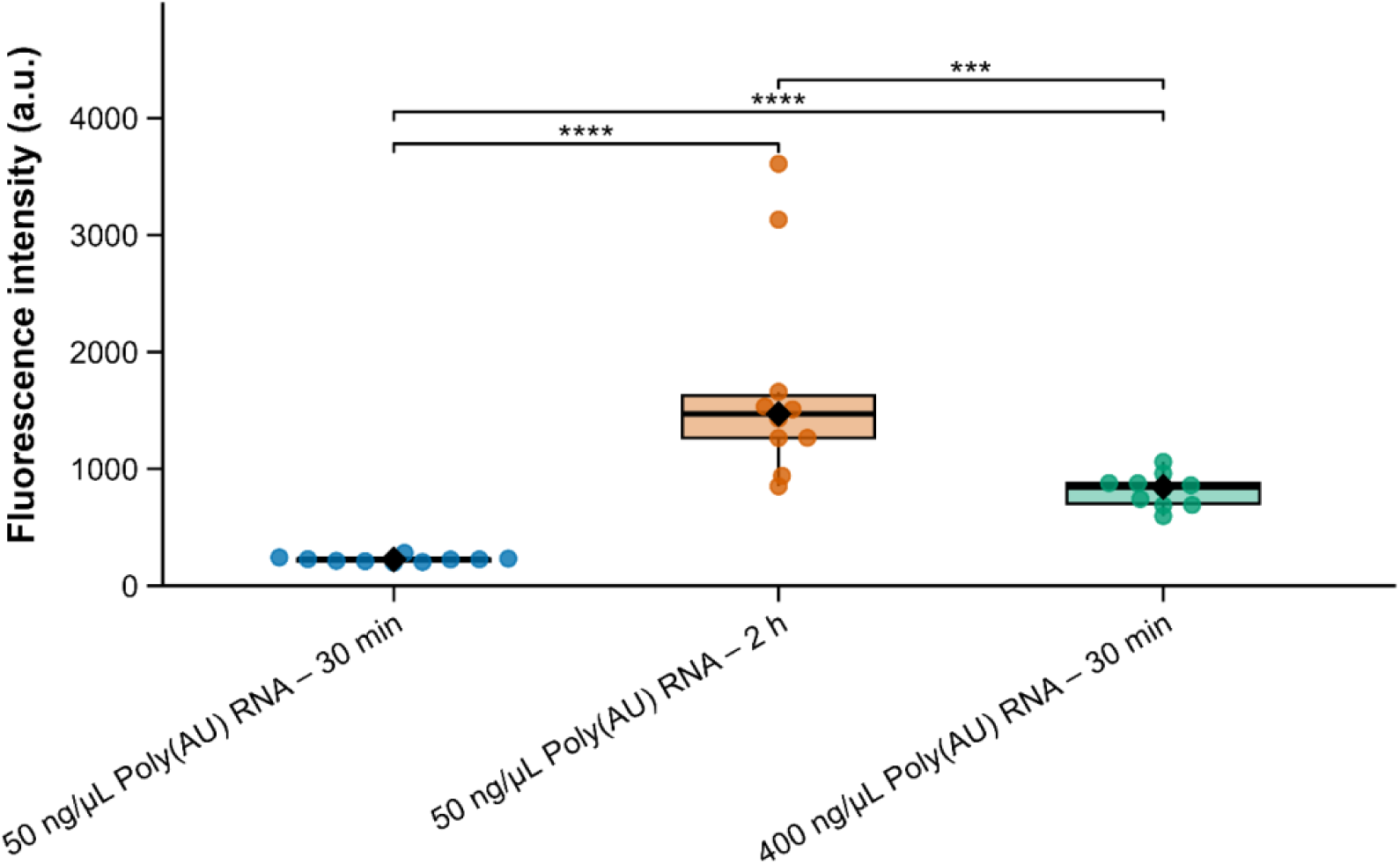
PSMα3 amyloid formation is enhanced by Poly (AU) RNA in a time-and concentration-dependent manner. The graphs show the calculated AmyTracker (AT630) fluorescence intensity of 100 µM PSMα3 incubated with Poly (AU) RNA under the indicated conditions, corresponding to the experiment shown in Figure 3. Boxplots show the interquartile range with the centre line indicating the median; black diamonds denote the mean. Individual points represent independent measurements. Statistical significance was assessed using a Kruskal–Wallis test followed by Dunn’s post hoc test with Bonferroni correction. *p<0.05, **p<0.01, ***p<0.001, ****p<0.0001.

**Figure S7.**
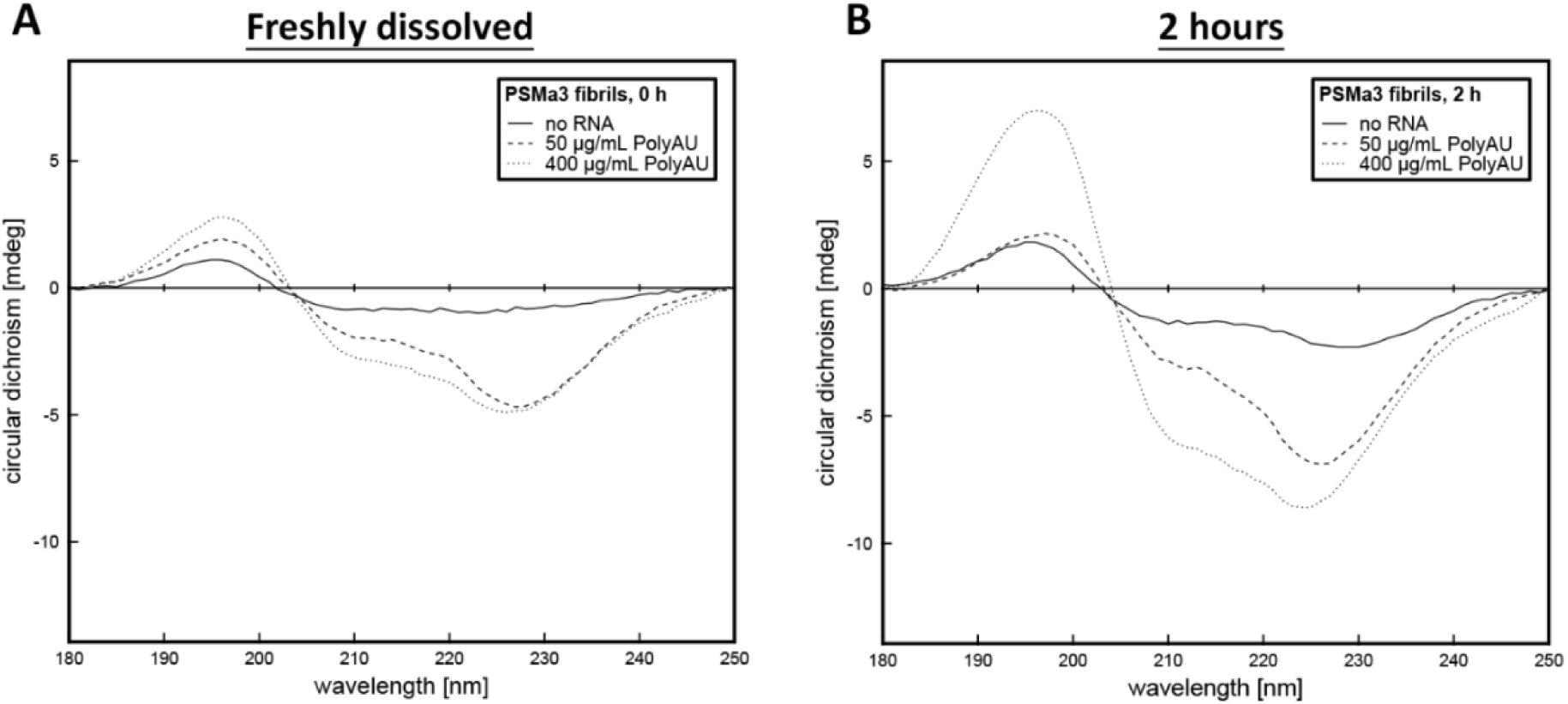
RNA enhances α-helical structural features of PSMα3 over time. Solid-state circular dichroism (ssCD) spectra of 100 µM PSMα3 incubated alone or with Poly (AU) RNA at concentrations of 50 ng/µL and 400 ng/µL. Spectra were collected immediately after preparation (**A**), and after 2 hours of incubation (**B**). Measurements were recorded in the far-UV range (180–250 nm) to assess changes in secondary structure under the indicated conditions.

**Figure S8.**
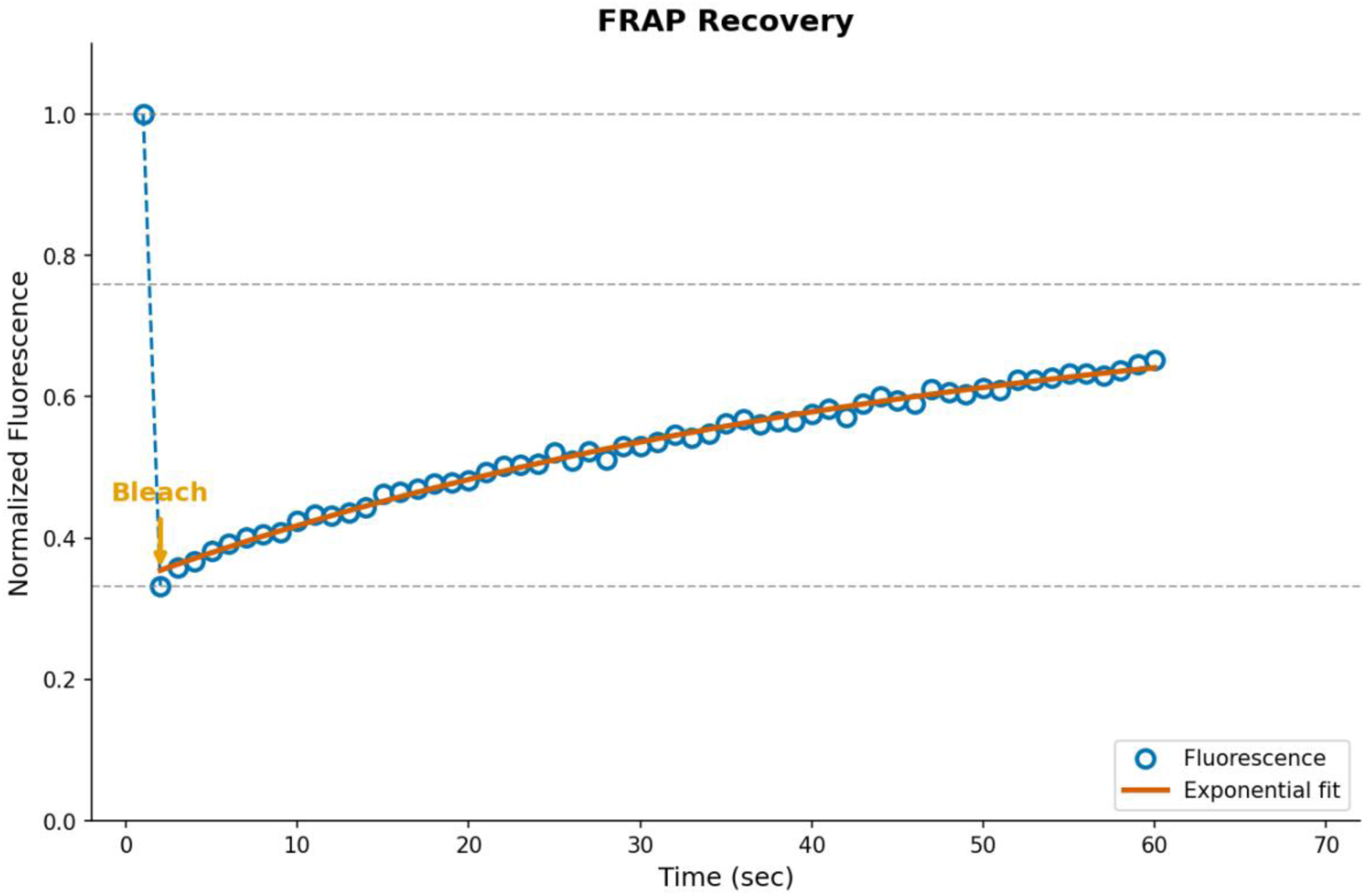
FRAP analysis of PSMα3 dynamics within the nucleolus of HeLa cells. Normalized FRAP of 20% FITC-PSMα3 and 80% unlabeled 20 µM PSMα3 measured within the nucleolus of HeLa cells, corresponding to the experiment shown in Figure 5. Blue circles represent fluorescence intensity values normalized to the pre-bleach signal, and the orange line shows a single-exponential fit to the recovery kinetics. The photobleaching event is indicated by the orange arrow. Recovery approached ∼64% within the 60 s acquisition window, suggesting the presence of both mobile and slowly exchanging populations. The recovery did not reach a clear plateau within the measured time frame, indicating the presence of a slowly exchanging or partially immobilized PSMα3 population within the nucleolus.

**Figure S9.**
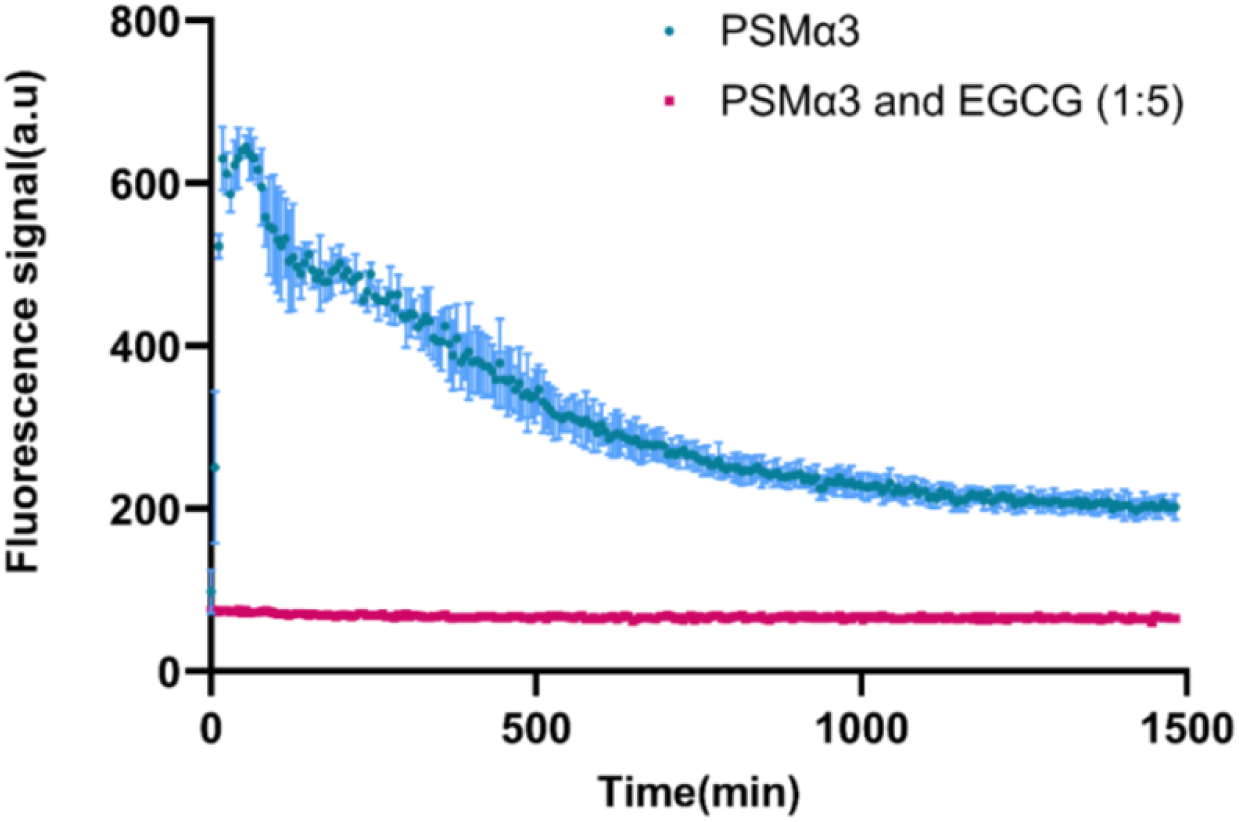
EGCG inhibits PSMα3 fibrillation. Fibrillation kinetics of 100 µM freshly dissolved PSMα3, monitored by Thioflavin-T (ThT) fluorescence. The data compares PSMα3 fibrillation in the absence (blue curve) and presence (red curve) of EGCG. The graph displays the mean fluorescence intensities from triplicate ThT measurements, with error bars representing the standard deviation.

**Figure S10.**
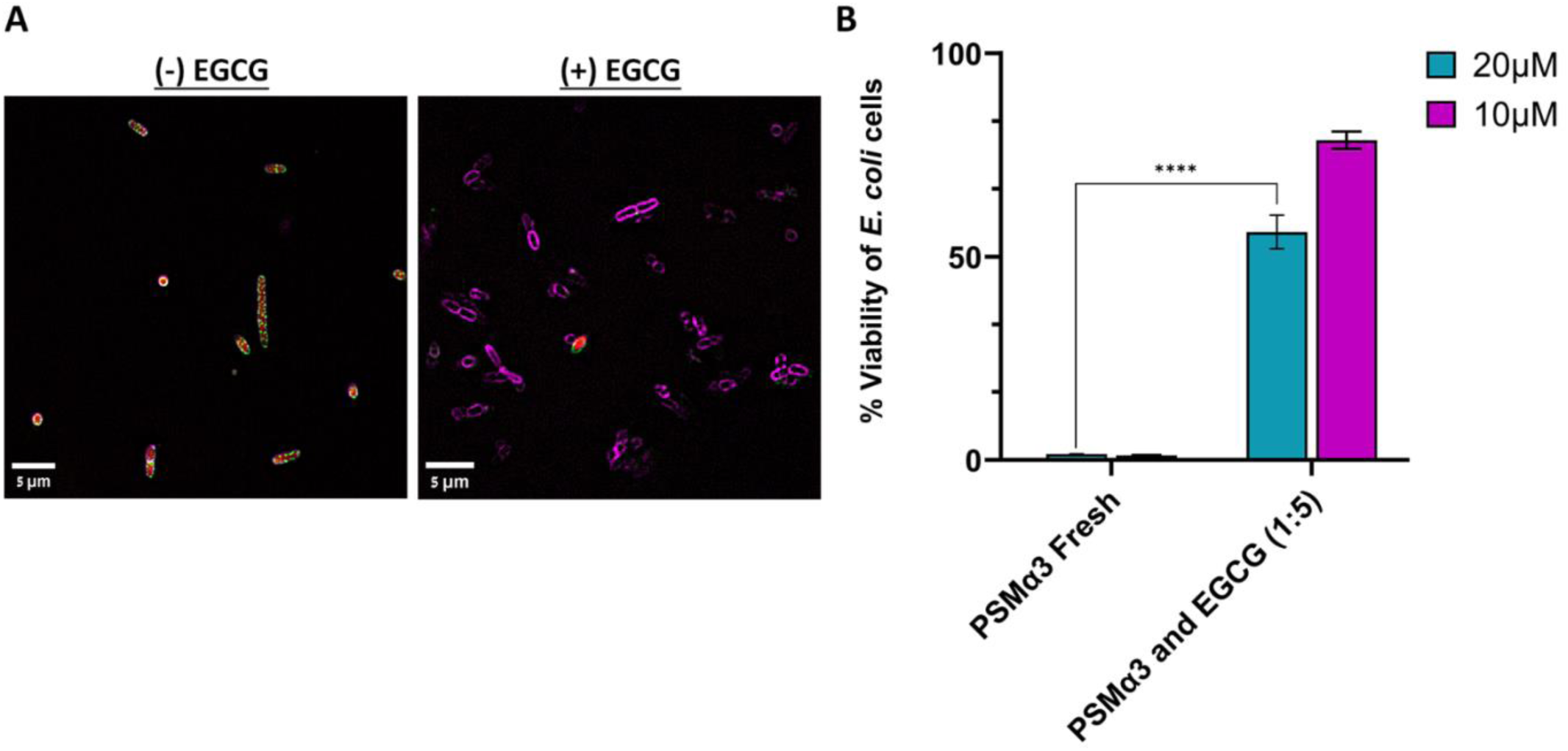
EGCG reduces the antimicrobial activity of PSMα3 against *E. coli*. **(A)** Super-resolution light microscopy images of *E. coli* treated with 20 μM of 20% FITC-PSMα3 (green) and 80% unlabeled PSMα3 in the absence (left) and presence (right) of EGCG at a 1:5 molar ratio. Scale bars: 5 μm. **(B)** Antimicrobial activity of PSMα3 against *E. coli*, evaluated using the PrestoBlue cell viability assay, with and without EGCG (1:5 molar ratio). The experiments were performed in at least three replicates and repeated across three independent days to ensure result reliability. Bacterial viability percentages were calculated as the mean of all replicates, with error bars representing the standard deviation. Statistical significance was determined using one-way ANOVA for normally distributed data in GraphPad Prism (version 11). Significance levels are indicated as follows: *p<0.05, **p<0.01, ***p<0.001, ****p<0.0001.

**Figure S11.**
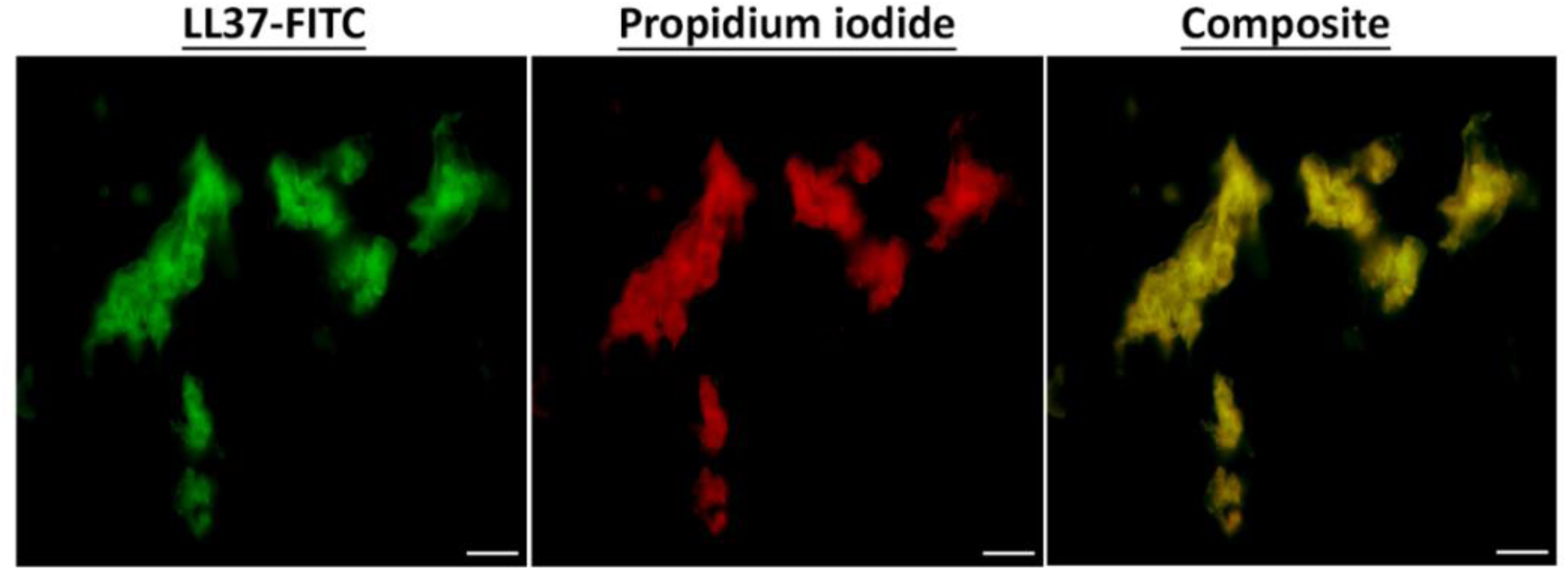
LL-37 undergoes aggregation with Poly (AU) RNA. Light microscopy images showing 20% FITC-labeled (green) and 80% unlabeled 100 μM LL-37 (with no thermal stress) forming aggregates in the presence of 100 ng/μL Poly (AU) RNA, stained with PI (red). The composite image (right) highlights the colocalization of LL-37 and RNA (yellow), indicating RNA-induced aggregation. Scale bars: 20 μm.

**Figure S12.**
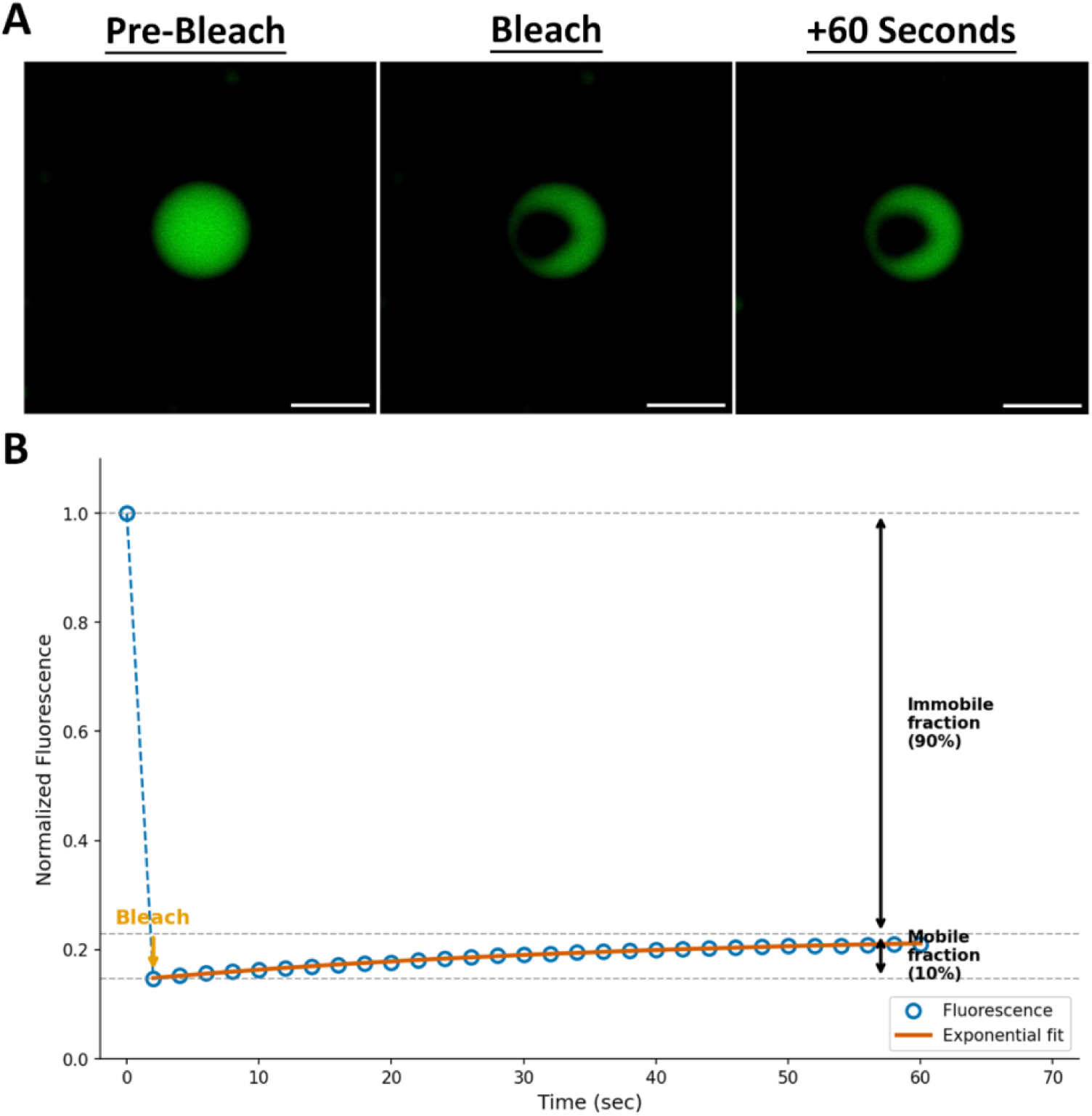
FRAP analysis of LL-37 – Poly (AU) RNA assemblies following heat shock. **(A)** Representative fluorescence images of 20% FITC-labeled (green) and 80% unlabeled 100 μM LL-37 assemblies formed in the presence of 100 ng/µL Poly (AU) RNA, shown before photobleaching (Pre-Bleach), immediately after photobleaching (Bleach), and 60 s after bleaching (+60 Seconds). **(B)** Normalized FRAP corresponding to the bleached region shown in (A). Blue circles represent fluorescence intensity normalized to pre-bleach levels, and the orange curve shows a single-exponential fit to the recovery data. The bleach event is indicated by the orange arrow. Recovery reaches a limited plateau corresponding to a mobile fraction of 10% and an immobile fraction of 90%. Horizontal dashed lines denote pre-bleach intensity (top), plateau recovery level (middle), and immediate post-bleach intensity (bottom).

**Figure S13.**
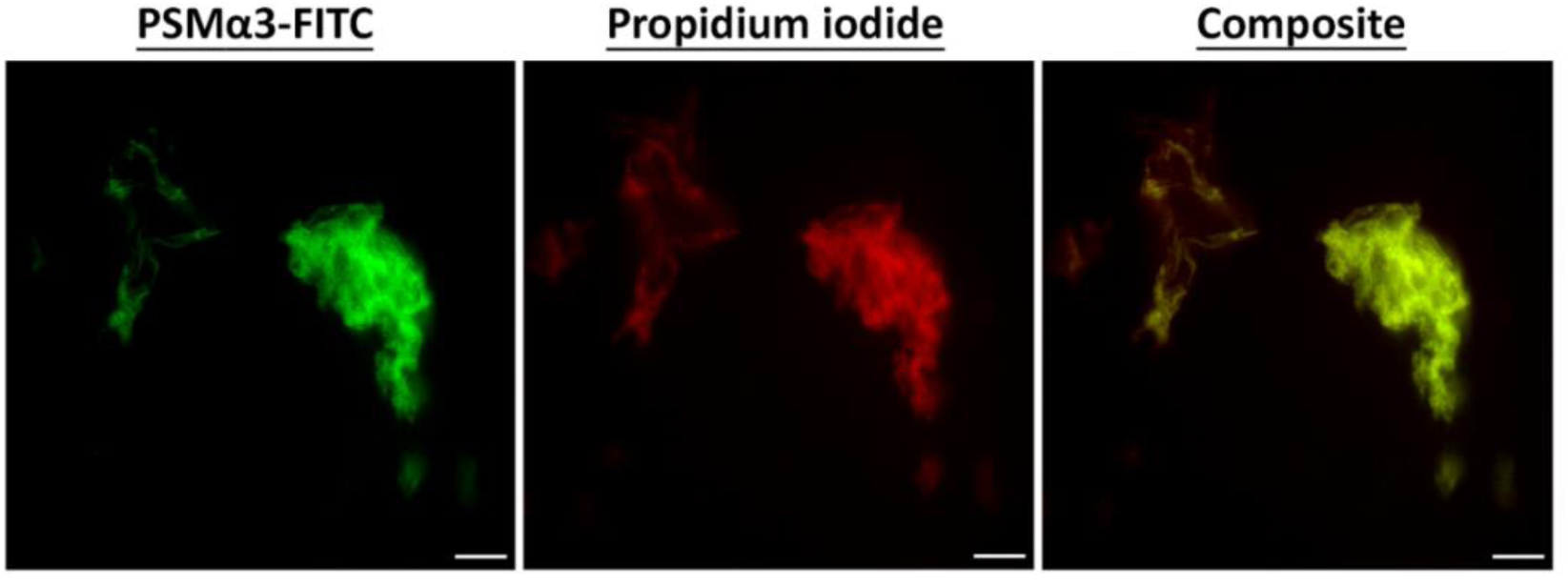
PSMα3 aggregates with Poly (AU) RNA after heat shock at physiological pH. Light microscopy images showing 20% FITC-PSMα3 (green) and 80% unlabeled 100 μM PSMα3 forming aggregates with 50 ng/μL Poly (AU) RNA, stained with PI (red), after heat shock at 65°C for 15 minutes in 50 mM HEPES, 150 mM NaCl, at physiological pH (7.4). The composite image (right) highlights the colocalization of PSMα3 and RNA (yellow), indicating RNA-induced aggregation under heat stress. Scale bars: 20 μm.

**Figure S14.**
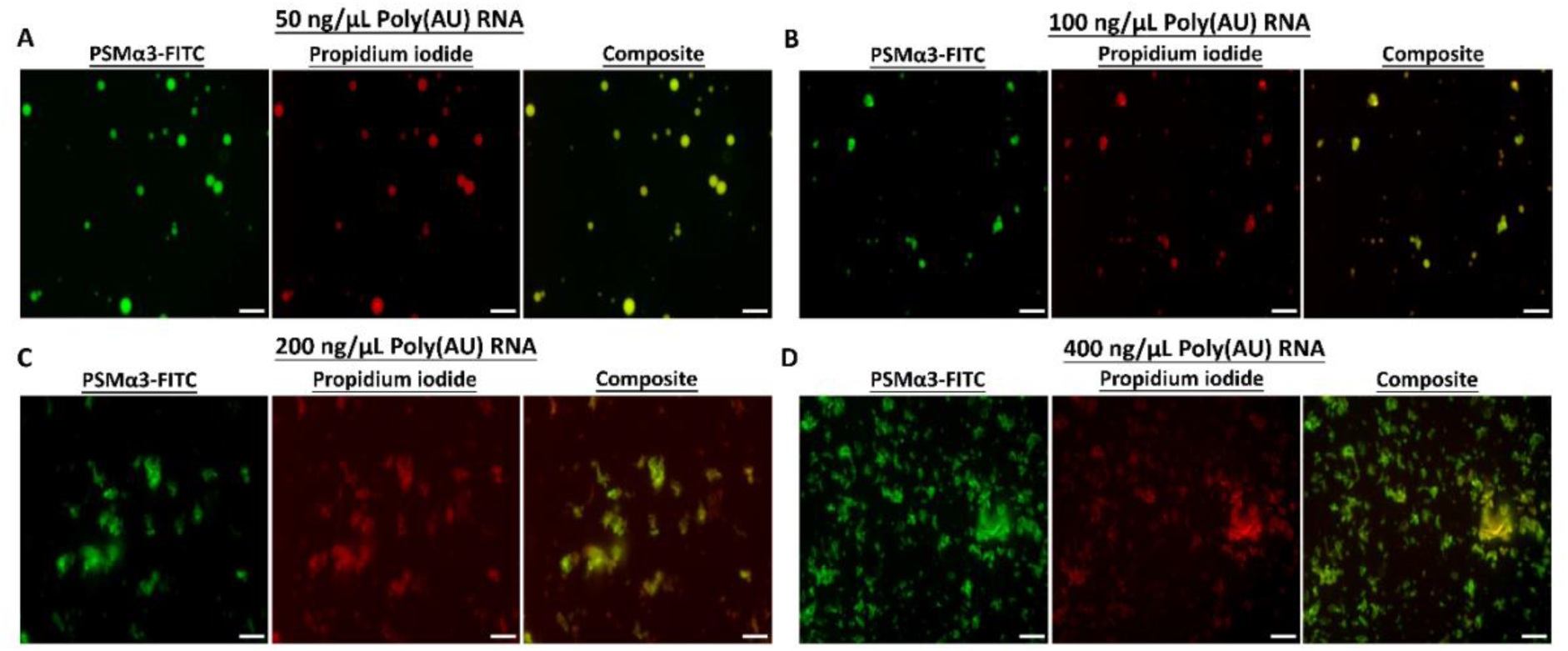
Effect of RNA concentration on PSMα3 phase separation and aggregation at pH 4 after heat shock. Fluorescence microscopy images showing the effects on 20% FITC-PSMα3 (green) and 80% unlabeled 100 μM PSMα3 in the presence of increasing concentrations of Poly (AU) RNA (labeled with PI, red) following heat shock at 65°C for 15 minutes at pH 4. At 50 ng/μL (**A**), 100 ng/μL (**B**), 200 ng/μL (**C**) and 400 ng/μL (**D**). At a low RNA concentration of 50 ng/μL (**A**), clear phase separation of PSMα3 was observed, as evidenced by the presence of well-defined, spherical droplets. As the RNA concentration increased to 100 ng/μL, the droplets began to lose their spherical structure and showed more morphological irregularities (**B**). At RNA concentrations of 200 ng/μL and 400 ng/μL, a distinct transition to aggregation with irregular, amorphous clusters was observed (**C** and **D**, respectively). The composite images revealed an overlap between the PSMα3 and RNA, indicating colocalizationduring both phase separation and amorphous cluster formation. Scale bars in all images represent 20 µm.

**Figure S15.**
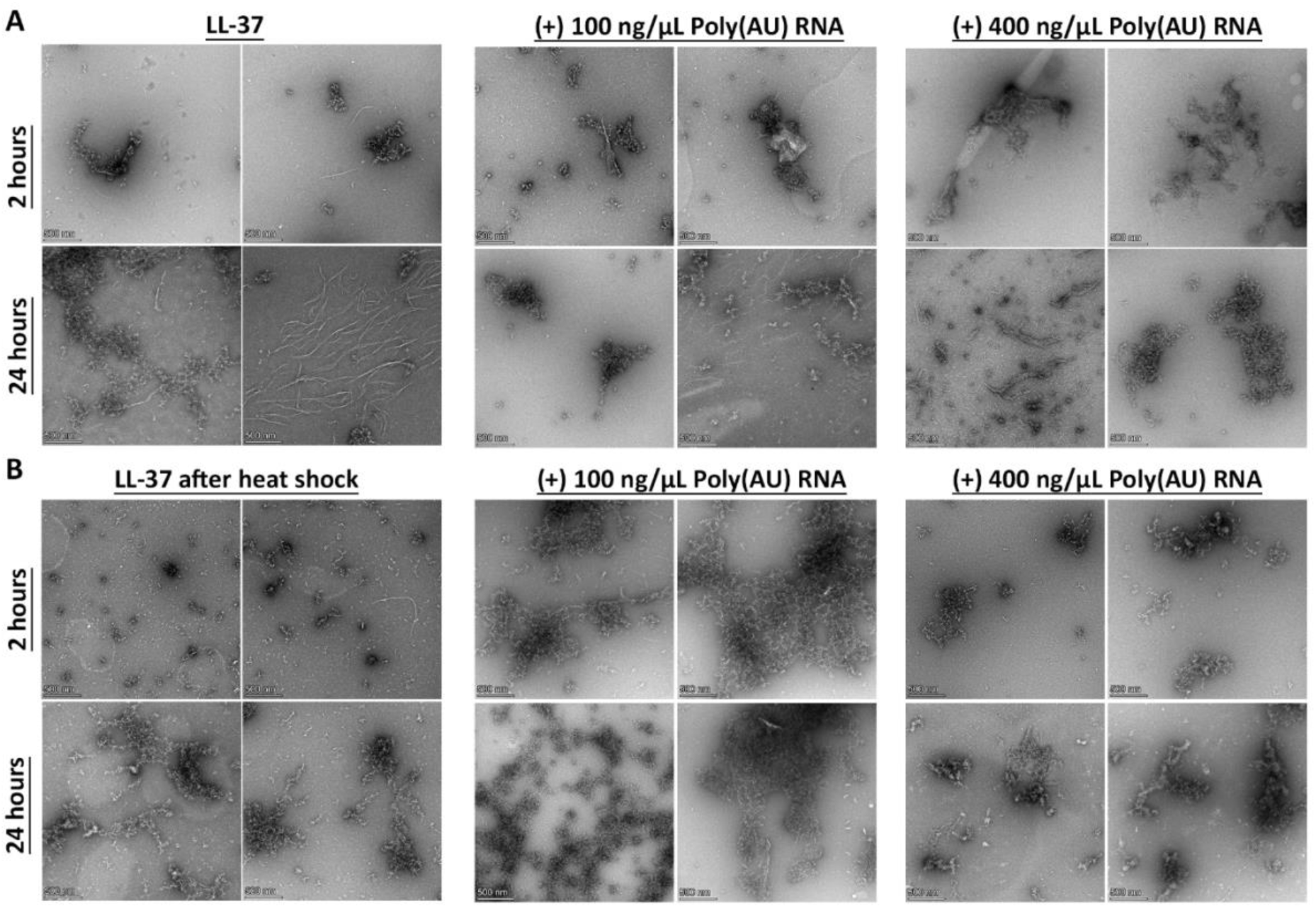
Poly (AU) RNA modulates LL-37 aggregation dynamics before and after heat shock. TEM micrographs of 100 µM LL-37 incubated alone or with 100 ng/µL or 400 ng/µL Poly (AU) RNA for 2 hours or 24 hours. **(A)** Samples imaged before heat shock at the indicated time points. **(B)** Samples subjected to a 65°C heat shock for 15 minutes, followed by further incubation for 2 or 24 hours before imaging.

**Movie S1. Live-cell imaging of PSMα3-induced membrane permeabilization in HeLa cells.** HeLa cells were incubated with 20 µM of 20% FITC-PSMα3 (green) and 80% unlabeled PSMα3 and monitored by time-lapse fluorescence microscopy. Nuclei were stained with Hoechst (blue). PI (red) was present in the imaging medium to detect loss of plasma membrane integrity. Time is shown in minutes: seconds (mm:ss) from peptide addition. Scale bar, 10 µm.

**Movie S2. Live-cell imaging of EGCG-modulated PSMα3 activity in HeLa cells.** HeLa cells were incubated with 20 µM of 20% FITC-PSMα3 (green) and 80% unlabeled PSMα3 in the presence of EGCG at a fivefold molar excess relative to PSMα3 (PSMα3: EGCG = 1:5) and monitored by time-lapse fluorescence microscopy. Nuclei were stained with Hoechst (blue). PI (red) was present in the imaging medium to detect loss of plasma membrane integrity. Time is shown in mm:ss from peptide addition. Scale bar, 10 µm.

